# NKG2D receptor ligands are cell surface biomarkers for injured murine and human nociceptive sensory neurons

**DOI:** 10.1101/2025.07.16.665122

**Authors:** Shuaiwei Wang, Allison M Barry, Yoon Kyung Lee, Naomi Young, Sang Wook Shim, Xinying Wang, Hyeongcheol Kim, Laura Stirling-Barros, Rafael González-Cano, Michael Costigan, Georgios Baskozos, Simon Rinaldi, David LH Bennett, Seog Bae Oh, Alexander J Davies

## Abstract

Nociceptors are primary afferent neurons that sense noxious stimuli. They can be activated by tissue injury as well as the accompanying local immune response. We have shown that following nerve injury in mice cytotoxic Natural Killer (NK) cells infiltrate the peripheral nerve and interact with stress-induced ligands of the activating receptor NKG2D (*Klrk1*). However, the diversity and specificity of NKG2D receptor ligands among sensory neuron subtypes, and translation of this mechanism to humans, remains unknown.

We used dorsal root ganglion (DRG) neurons cultured from C57BL/6J mice of both sexes with fluorescently-labelled sensory neuron lineages (*Scn10a*, *Mrgprd, Calca, Trpv1*, *Th*, *Thy1*), as well as human induced pluripotent stem cell derived (hiPSCd)-sensory neurons after laser ablation, as *in vitro* models of axonal injury. We assessed expression of NKG2D ligands by quantitative polymerase chain reaction (PCR) corroborated by publicly available RNA sequencing datasets and validated with single-cell PCR. Recombinant NKG2D receptor proteins in live cell-based assays were used to reveal the subcellular membrane localisation of NKG2D ligands with quantification by a semi-automated image analysis. Functional interactions between human NK cells and sensory neurons were confirmed with co-cultures in microfluidic devices.

We show that NKG2D ligands are expressed exclusively in unmyelinated DRG neurons after injury. NKG2D-receptors bound to puncta along distal neurites of injured axons enriched predominantly in *Mrgprd*-expressing non-peptidergic nociceptors. We observed low-level binding of human NKG2D-receptors to neurites of hiPSCd sensory neurons that increased after axonal laser ablation. Degeneration of hiPSCd sensory neurons neurites by interleukin (IL-2) primed human NK cells was prevented by an NKG2D blocking antibody.

The induction and enrichment of functional NKG2D receptor ligands selectively on pathological nerve fibres could aid the diagnosis of peripheral neuropathy in chronic pain conditions, and sheds new light on the potential role of nociceptive neurons in regulating the local tissue immune microenvironment.

## Introduction

Nociceptors are a specialized subset of primary afferent (somatosensory) neurons that respond to noxious stimuli [1] and are the first part of a neuronal pathway sending signals to brain that are perceived as pain. After injury or disease, peripheral nerves may undergo Wallerian degeneration followed by regeneration [2], as well as alterations to their biochemical and functional properties that are thought to contribute to peripheral sensitization and neuropathic pain [3]. Despite substantial evidence from preclinical studies, many novel, neuronally-targeted treatments for neuropathic pain have failed in clinical trials [4,5]. The current situation is reinforced by a recent systematic review and meta-analysis of clinical trial data on pharmacotherapeutic interventions which found a persistent reliance on existing drugs such as tricyclic anti-depressants, serotonin and norepinephrine reuptake inhibitors and α2δ-ligands as first list treatments for neuropathic pain [6], guidance which has not changed significantly in the last 10 years [7]. Understanding the broader changes in nociceptor form and function after nerve injury is therefore crucial for uncovering the mechanisms of neuropathic pain and developing effective treatments [8].

Recent developments in single cell sequencing technologies have greatly deepened our understanding of sensory neuron heterogeneity [9], as well as key differences between species [10,11]. However, our understanding of the pathological changes that occur to DRG neurons is hindered by the loss of transcriptional identity after nerve injury [12]. The resulting changes in expression of sensory neuron-specific drug targets for pain, such as Na_v_1.8 [13], are therefore likely to limit the re-purposing of promising pharmacological treatments for acute pain [14] to chronic neuropathic pain. Knowledge of an injury-specific biomarker, particular one expressed at the cell surface, would therefore greatly enhance our ability to identify and target pathological sensory nerves for potential therapeutic intervention.

In addition to intrinsic neuronal changes, the immune system has a critical role in peripheral sensitization and pain following nerve injury [15], with bidirectional neuro-immune crosstalk between nociceptors and immune cells influencing the outcomes of pain and inflammation [16,17]. Our previous work has highlighted the role of cytotoxic natural killer (NK) cells in the immune response to nerve injury and neuropathic pain resolution [18], suggesting immunotherapies targeting innate cytotoxic cells as a novel potential strategy for neuropathic pain treatment [19].

NK cells, a key component of innate immune system and member of the innate lymphoid cell (ILC) family, perform immuno-surveillance for transformed, virus-infected and stress cells within the body, as well as regulate adaptive immune responses [20]. Among NK cells’ direct functions are the recognition of stress-induced molecules, or “altered-self”, through their activating receptors [21,22]. The archetypal activating receptor Natural Killer group 2D (NKG2D), encoded by the gene *Klrk1* in mice (*KLRK1* in humans) [23], serves as an evolutionarily conserved a “master switch” regulating NK cell activation in both species [24,25]. In mice, NKG2D ligands are encoded by the *Raet1* gene family consisting of five gene isoforms: α, β, γ, δ and ε [26,27]. Other known NKG2D ligands include the multi-gene family histocompatibility 60 (*H60)* a-c [28] and murine UL16-binding protein-like transcript 1 *Mult1 (Ulbp1)* [29], which also contribute to NK cell-mediated cytotoxicity (Lanier, 2015). In humans, NKG2D ligands include MHC class I polypeptide-related sequence A (MICA) and MICB, and the UL16-binding protein family: ULBP1 (RAET1I), ULBP2 (RAET1H), ULBP3 (RAET1N), ULBP4 (RAET1E), ULBP5 (RAET1G), and ULBP6 (RAET1L) [30,31]. Engagement of the NKG2D receptor with ligands expressed at the cell surface as transmembrane or glycophosphatidylinositol (GPI]-linked membrane proteins may render the target susceptible to cell-mediated cytotoxicity by NK or CD8+ T cells, and as well as subsets of NKT and γδ T cells [32,33].

We have previously demonstrated that genes for the *Raet1* family of ligands are upregulated within dorsal root ganglia (DRG) following peripheral nerve injury in mice [18]. However, the diversity of the changes in NKG2D ligand expression across sensory neuron subtypes, and therefore the cellular specificity of targeting by NK cells after nerve injury, remains undefined. In this study, we aimed to identify the subpopulation of sensory neurons expressing activating ligands for NKG2D after injury, as well as examine the potential for translation of this cytotoxic neuro-immune interaction from mice to humans.

## METHODS AND MATERIALS

This study is reported in accordance with the RIVER (Reporting In Vitro Experiments Responsibly) recommendations for *in vitro* experiments [34] and ARRIVE (Animal Research: Reporting of In Vivo Experiments) guidelines for animal experiments [35].

### Ethical approvals

Human NK cells were isolated from the peripheral blood of healthy donors collected by the UK National Health Service (NHS) Blood and Transplant service and distributed by Non-Clinical Issue (NCI) with the approval of the University of Oxford Medical Sciences Interdivisional Research Ethics Committee (MS IDREC) (Reference: R70042/RE002) and stored under a Human Tissue Authority site licence (HTA_12217; Project 00122). Peripheral blood was also collected from healthy donor volunteers after informed consent with the approval of the South Central–Oxford A Research Ethics Committee (14/SC/0280). All nerve injury and capsaicin administration procedures were approved by the Institutional Animal Care and Use Committee (IACUC) at Seoul National University (SNU-121011-1) in Korea and Boston Children’s Hospital (15-04-2928R and 16-01-3080R) in the USA. Tamoxifen dosing in inducible *cre* lines was performed under a UK Home Office Project Licence (P1DBEBAB9). Animals were killed according to Schedule 1 of the UK Home Office (Scientific Procedures) Act (1986).

### Animals

All mice were group-housed in individually ventilated cages with free access to food and water, in humidity and temperature-controlled rooms with a 12hr light-dark cycle (lights on 07.00am), in a pathogen free facility. For collecting DRG tissues, all the mice were culled using Schedule 1 by exposure to a rising concentration of carbon dioxide followed by cervical dislocation. *Calca^creERT2^*(Calca^tm1.1(cre/ERT2)Ptch^) were kindly gifted by Prof. Pao-Tien Chuang, *Nav1.8^cre^* (*Scn10a^tm2(cre)Jnw^*) mice were kindly gifted by Prof. John Wood, and Thy-1 YFP-16 (B6;Cg-Tg(Thy1)-^(YFP)16Jrs/J^) mice (RRID:IMSR_JAX:003709) were kindly gifted by Prof. Pilhan Kim. *Mrgprd^creERT2^* mice (*Mrgprd*^tm1.1(cre/ERT2)Wql/J^) (RRID:IMSR_JAX:031286), *Th^creERT2^* mice (*Th*^tm1.1(cre/ERT2)Ddg/J^) (RRID:IMSR_JAX:025614) were kindly gifted by Prof. David Ginty, *Trpv1^cre^* (*Trpv1^tm1(cre)Bbm^/J*) (RRID: IMSR_JAX:017769), *R26R^DTA^* (*B6;129-Gt(ROSA)26Sor^tm1(DTA)Mrc^/J*) (RRID:IMSR_JAX:010527) and Ai14 (B6.Cg-Gt(ROSA)26Sor^tm14(CAG-tdTomato)Hze^/J) (RRID:IMSR_JAX:007914) mice were purchased from Jackson Labs and have been previously described [36–40]. Cre-dependent reporter lines were maintained with heterozygote inbreeding. Heterozygote cre-dependent reporter mice were crossed with *DTA* or Ai14 ‘TdTomato’ reporter mice generate conditionally ablate or fluorescently label of selected sensory neuronal lines, respectively. Double-heterozygous mice were used for experiments. Male and female C57BL/6J mice were purchased from the Envigo (Inotiv) in the UK, Dae Han Bio Link (Taconic) in Korea, or Jackson Laboratories (Jax) in the US, and were used 8-12 weeks of age. Transgenic mice were used aged 12-40 weeks of age. Owing to unpredictable breeding patterns, and to maximise the use of tissues, *Mrgprd^creERT2^* mice used in this study had received spared nerve injury to the left sciatic nerve 30 days prior to cell culture for other experiments; bilateral L3-5 DRG were therefore excluded from the cell culture used for receptor-binding analysis.

### Tamoxifen administration

All cre/ERT2 expressing mice were dosed 5x (daily) with 75 mg/kg Tamoxifen (Sigma-Aldrich) in adulthood (8+ weeks), as reported previously [41]. Tamoxifen stocks were freshly prepared by dissolving at 20 mg/ml in corn oil via sonification. All animals were dosed i.p. and health statuses were monitored daily for the duration of the dosing regimen.

### Peripheral nerve injury

For L5 spinal nerve transection (L5x) injury mice were placed under isoflurane anaesthesia by inhalation (3% induction, maintained 1-2% in 99% O_2_ at 1-2 L/min), the dorsal lumbar region was shaved, treated with an iodine solution (Potadine) and a unilateral incision made parallel to the L6 vertebrate. Under a x20 dissection microscope illuminated by a cold light source the musculature was parted by blunt forceps dissection to reveal the L6 transverse process, which was then cut and removed. The L5 spinal nerve, which runs immediately below the L6 process, was carefully freed of connective tissue and cut with fine spring scissors; 1 mm of the nerve was removed to prevent nerve regeneration. The wound as irrigated with sterile saline and closed in two layers with 6-0 silk sutures (Ailee, Korea) and 9mm skin clips (MikRon Precision, CA, USA) [18]. For spared nerve injury (SNI), mice were anesthetized with isoflurane (2%–4%) at 9 weeks and SNI surgery performed; the tibial and common peroneal branches of the sciatic nerve were tightly ligated with a silk suture and transected distally, whereas the sural nerve was left intact [42]. Mice were placed in a warm, darkened cage to recover from surgery and monitored daily for signs of malaise, piloerection, weight loss or autotomy. No post-surgical analgesia was provided.

### TRPV1-expressing neuron ablation

TRPV1-expressing neurons were ablated with resiniferatoxin (RTx) as previously described [43]. Briefly, under isoflurane anaesthesia 3-4 week old male mice were injected i.p. with RTx at 50 µg/kg and 150 µg/kg or equivalent volume of vehicle (10% ethanol, 10% tween-80 in sterile saline) over two consecutive days. Mice were maintained under anaesthesia (0.5% isoflurane in 100% O_2_ at 1-2 L/min) for 2 h on a warm-pad before recovery in a warmed cage. Four weeks after RTx or vehicle treatment mice were tested for TRPV1+ neuron depletion by applying 10 µl capsaicin (0.01% in sterile saline) to one eye and counting the number of unilateral eye wipes in a 1 min period. Video analysis was performed by an observer blinded to the treatment of the mice.

### DRG neuron culture

Dorsal root ganglion (DRG) neurons were cultured as previously described [18,44]. Mice were killed by rising concentration of carbon dioxide and death confirmed by cervical dislocation. The spinal column was rapidly dissected and placed in ice cold Ca^2+^ and Mg^2+^-free Hank’s Buffer Saline Solution (Gibco, 14065-049) supplemented with 20mM HEPES). Individual DRG were dissected and trimmed of nerve roots and digested 60 min in collagenase A (1 mg/ml) (Cat. 10103578001, Roche) and dispase II (2.4U/ml) (Cat. 04942078001, Roche) at 37°C. Additional digestion was carried out for 5-7 min in trypsin (0.25%) in Dubellco’s Modified Eagle Medium (DMEM) (Cat. 41965-039, Gibco) and stopped with a trypsin inhibitor (2.5 mg/ml) (Cat. T9003, Sigma) in PBS followed by washing in DMEM containing 10% foetal bovine serum (FBS) (Life Technologies). DRG were dissociated by trituration with a fire-polished glassed pipette in DMEM containing DNase I (125 U/ml) (Cat. 58409700, Roche) and centrifuged 10 min at 200 g on a layer of bovine serum albumin (BSA) Cat. A7248, Sigma) diluted to 15% in DMEM, before re-suspension in neurobasal medium (Cat. 12348017, Life Technologies) with B27 supplement (Cat. 17504-004, Life Technologies), L-glutamine (1 mM), penicillin (100 U/ml) and streptomycin (100 U/ml) (Cat. 15140-122, Life Technologies) supplemented with nerve growth factor (NGF 2.5S) at 50 ng/ml. 10^3^ DRG neurons were plated on 13 mm diameter glass coverslips previously coated with poly-D-lysine (PDL) (10 μg/ml) (Cat. P6406, Sigma) and laminin (10 μg/ml) (Cat. L2020, Sigma) and maintained in culture at 37°C, 5% CO_2_, for up to 3 days prior to immunolabeling.

### Microfluidic device preparation

Microfluidic devices created from polydimethylsiloxane (PDMS) (Cat. 63416.5S, Silicone Sylgard 184, Scientific Laboratory Supplies Limited). Slygard 184 silicon base (density approximately 1g/ml) and curing agent were mixed in a 10:1 ratio. The mixture was rested for 15 min to remove bubbles before pouring into resin master moulds, which were placed in oven for at least one hour at 60-65°C to set. After removing from the resin master moulds, PDMS devices were trimmed and four reservoirs were cut using an 8 mm diameter skin punch biopsy tool (Cat. BP-80F, Selles Medical Ltd). Two of the reservoirs on the neurite side were enlarged to make a single continuous reservoir. The length of microfluidic channels was 150 µm. The trimmed PDMS devices were adhered to low profile glass-bottom dishes (Cat. HBST-5040, Willco wells) by pre-exposure in a low-pressure plasma cleaner (Femto, Diener Electronic GmbH) 2x 30s at 35% power. Approximately 150 µl sterile distilled water was immediately added in one of the smaller reservoirs to maintain patency of the microfluidic channels, and devices were kept at 4 °C before use.

### Human induced pluripotent stem cell derived (hiPSCd) sensory neuron cultures

hiPSCs from healthy control donors were obtained via the University of Oxford StemBANCC consortium (**Supp. Table 1**). hiPSC lines were characterised by probe-based karyotyping (Karyostat Assay, Applied Biosystems) and confirmed free from mycoplasma (MycoStrip, InvivoGen). hiPSC were maintained in 6-well culture plates coated with hESC-qualified LDEV-free Matrigel (Cat. 354277, Corning) with daily changes of mTeSR1 media (Cat. 85850, StemCell Technologies). Cells were routinely passaged by EDTA dissociation (500 µM in PBS, 5 min) when cells became 70-80% confluent. hiPSCs were differentiated into sensory neurons in 6 well plates using a combination of small-molecule dual SMAD (mothers against decapentaplegic family transcription factor) inhibitors SB431542 (10 µM) (Cat. 1614, Tocris) and LDN-193189 (100 nM) (Cat. 6053, Tocris) and Wnt pathway activators CHIR99021 (3 µM) (Cat. 4423, Tocris), SU5402 (10 µM) (Cat. SML0443, Sigma), and DAPT (10 µM) (Cat. 2634, Tocris) [45,46]. Briefly, hiPSC media was changed to Mouse Fibroblast-conditioned media (MEF) (Cat. AR005, R&D Systems) supplemented with Fibroblast growth factor (FGF)-2 (10 ng/ml) for one day (Day −1). The next day (Day 0) differentiation began with the addition of a series of small molecule inhibitors added to Knock-out Serum Replacement (KSR) media: Knockout DMEM (Cat. 10829018, life Technologies) containing 15% knockout-serum replacement (Cat 10828010, Life Technologies), 1% Glutamax, 1% non-essential amino acids, 1% antibiotic/antimycotic, and β-mercaptoethanol (100 μM). Media was changed daily (2 ml per well), during which time KSR media was gradually replaced with N2 ‘complete’ media (neurobasal medium supplemented with N2 (Cat. 17502-048), B-27 minus vitamin A (Cat. 12587-010, Gibco), Glutamax (Cat. 35050-038, Gibco) and 1x antibiotic-antimycotic (Anti-anti) (Cat. 15240-062, Gibco) plus recombinant human β-NGF (rhNGF) (Cat. 450-01, Peprotech), NT3 (Cat. 450-03, Peprotech), GDNF (Cat. 450-10, Peprotech), and BDNF (Cat. PHC7074, Life Technologies) at 25 ng/ml each), according to published protocols [46,47]. On day 11 or 12 of differentiation, cells were washed with 1 ml per well of PBS, followed by knockout DMEM containing DNase I (100 U/ml) and incubated 5 min at 37°C, 5% CO_2_. The knockout DMEM was then replaced with 1 ml TrypLE dissociation reagent and incubated for a further 5 min at 37°C, 5% CO_2._ before dissociation with fire-polished glass pipette. Cells were transferred to PBS in 15 ml tubes and centrifuged 5 min, 500 g to pellet. Supernatant was removed and cells were resuspended in N2 ‘complete’ media supplemented with CHIR99021 (3 µM) and Rho kinase inhibitor, Y-27632 (10 µM) (Cat. 1254, Tocris). Cells were counted, suspension at 4×10^6^ cells per ml in N2 completed neuronal media (above), diluted 1:1 with chilled 2X freeze media (20% DMSO, 80% charcoal-stripped FBS) and immediately transfer cell suspension to cryovials (2×10^6^ cells per ml per vial).

Thawed precursor neurons were seeded either onto 13 mm diameter glass coverslips (approximately 20,000 cells per coverslip), 24-well glass bottom plates (60,000 cells per well) (Cat. 662892, Greiner-One Bio), or microfluidic devices (50,000 cells per device) previously coated with poly-D-lysine (PDL) (10 μg/ml) followed by reduced growth-factor Matrigel (Cat. 354277, Corning) or Geltrex (LDEV-Free, hESC-Qualified) (Cat. A1413302, ThermoFisher). Sensory neuron precursors were maintained in N2 ‘complete’ media supplemented with CHIR99021 (3 µM) and Rho kinase inhibitor, Y-27632 (10 µM) for the first two days after seeding, after which the neuronal media was supplemented with cytosine arabinoside (Ara-C) (0.5-1 µM) (Cat. C1768, Sigma) to remove proliferating, non-neuronal cells. Neurons were matured in N2 ‘complete’ media in an incubator at 37°C, 5% CO_2_ for at least 4 weeks (40 days after differentiation), with media changes twice per week.

### hiPSCd-sensory neuron axon ablation for NKG2D receptor labelling

Cryopreserved sensory neuron precursors were thawed and seeded (60,000 cells/well) onto 24-well glass bottom plates (Cat. 662892, Greiner Bio-One Ltd) coated with PDL (10 µg/ml) and Geltrex and cultured in N2 ‘complete’ media. After 4 weeks the axons of hiPSCd-sensory neurons were ablated in a line in the centre of each well by a 355 nm UV laser (Rapp OptoElectronic, model: DPSL-355/42/CLS2) 3% laser power controlled by SysCon software under visual control via a spinning disc confocal microscope (iXplore SpinSR10, Olympus). Cultures were maintained up to one week to allow regenerated neurites to be assessed in subsequent experiments.

### NKG2D receptor binding and immunolabelling

Recombinant murine NKG2D-Fc chimeric receptor (Cat. 139-NK, R&D Systems; lot numbers: FRP0218081, FRP0222021, FRP0223061, FRP0224041), recombinant human NKG2D-Fc chimeric receptor (Cat: 1299-NK, R&D Systems; lot number: FVV0618021) and recombinant human IgG1 Fc control (Cat. 110-HG, R&D Systems, lot number: EAX0619041) were prepared by suspension in distilled water at 1 mg/ml for 15 min at room temperature (RT) and further diluted to 100 µg/ml in phosphate buffered saline (PBS) containing 0.1% BSA for storage at −80°C. Some lot-lot variation was observed with murine NKG2D-Fc proteins; lot number FRP0220051 was found in trials to bind poorly to DRG neuron cultures and was therefore not used for experiments. Receptor proteins were diluted to 2 µg/ml in neurobasal media containing 1% BSA and applied to live DRG neurons on coverslips for 1h at 37°C. Cells were gently washed three times with PBS and fixed with 4% PFA in PBS for 30 min at room temperature (RT) before washing with PBS followed by two washed with DMEM containing 20 mM HEPES (DMEM/HEPES). To immunolabel Fc-conjugated receptor proteins, coverslips were treated with either Alexa 488-conjugated goat anti-human IgG (1:750) (Cat. A-11013, Thermo Fisher Scientific, RRID: AB_2534080) or Cy3-conjugated goat anti-human IgG (1:750) (Cat. 109-165-170, RRID: AB_2810895) in DMEM/HEPES and 1% BSA for 1h at RT. Coverslips were washed with DMEM/HEPES followed by PBS. Cells were blocked and permeabilised by incubation with 5% normal goat serum (NGS) and 0.1% triton-X in PBS for 1h at RT and incubated with rabbit anti-β-tubulin III (1:2000) (Cat. T2200, Sigma-Aldrich; RRID: AB_262133) overnight at 4°C. The next day, after washing in PBS coverslips were treated with either Alexa-546 conjugated goat anti-rabbit IgG (1:1000) (Cat. A-11035, Thermo Fisher Scientific, RRID: AB_2534093), Pacific Blue-conjugated goat anti-rabbit IgG (1:1000) (Cat. P-10994, Thermo Fisher Scientific, RRID: AB_2539814), Alexa Fluor 647 conjugated donkey anti-rabbit IgG (1:1000) (Cat. A-31573, Thermo Fisher Scientific, RRID: AB_2536183) or Alexa Fluor 647 conjugated goat anti-rabbit IgG (1:1000) (Cat. A-21244, Thermo Fisher Scientific, RRID: AB_2535812) for 1h at RT (see **Supp. Table 2**). Coverslips were washed in PBS, mounted on glass slides with antifade mounting medium (Vectorshield Plus, Cat. H-1000, Vector Laboratories) and stored at −20°C until imaging; microfluidic devices were flooded with PBS and stored at 4°C.

### Plasmid amplification and purification

Commercial plasmids containing gene inserts encoding murine RAE-1-ε (untagged) or green fluorescent protein (GFP) control under the control of the CMV promoter (see **Supp. Table 3**) were transferred to competent *E. coli* (NEB® Stable, Cat. c3040i, New England Biolabs) by heat-shock treatment. Briefly, 100 ng/μl plasmid DNA was added to 15 ul *E. coli, i*ncubated on ice for 30 minutes, followed by heat-shock at 42°C for 30 s, and immediately returned to ice for a further 5 min. SOC outgrowth media (250 μl) (Cat. B9020S, New England Biolabs) was added to the plasmid/*E. coli* mixture and incubated at 37°C with shaking (250 rpm) for 1 h. 10 to 20 μl of the starter culture was then spread onto plates prepared from LB agar (3.2%) (Cat. L3027, Sigma-Aldrich) containing ampicillin (100 µg/ml) (Cat. A5354, Sigma-Aldrich) and incubated overnight at 37°C. The next day a single colony was picked from each plate using a pipette tip and added to 5 ml of broth with ampicillin (100 µg/ml), followed by incubation for 4h in a shaking incubator (250 rpm) at 37°C to form a starter culture. Subsequently, 1 ml of the starter culture was transferred to 250 ml of LB broth (2.0%) (Cat. L3522, Sigma-Aldrich) with ampicillin (100 µg/ml) in sterile a 1-liter flask and incubated overnight at 37°C in a shaking incubator (250 rpm).

Plasmid DNA was purified using a PureLink™ HiPure Maxiprep kit (Cat. K210006, ThermoFisher Scientific). Briefly, cell cultures were pelleted by centrifugation (4600 g, 15 min at RT) to remove supernatant. Pellets were resuspended in suspension buffer, followed by an equal volume of lysis buffer, mixed by inversion and incubated 3 min at RT. Lysates were transferred to a filter column to remove cell debris. The lysate was clarified with binding buffer (10 ml) and passed through a purification column by centrifugation (500g, 2 min) followed by two rounds of wash buffer (800 µl, 17,000g, 1 min). DNA was eluted with 100 µl elution buffer; concentration and purity were confirmed on a spectrophotometer (1-3 ng/µl; 260/230 nm ratio >2.0) (Nanodrop). DNA inserts were confirmed by long-read sequencing (Nanopore-30, Source Bioscience).

### Heterologous expression of mouse Raet1e in HEK293T cells

HEK293T cells were seeded on coverslips previously coated with PDL (10 µg/ml) in 500 μl DMEM (Cat. 41965-039, Gibco) including 10% heat-inactivated FBS (Cat. 16140071, Life Technologies) in a 24-well plate (5×10^4^ cells per well) and incubated overnight at 37°C, 5% CO_2_. After approximately 48h (2 days) in culture, HEK cells were transfected with plasmid DNA using polyethylenimine (PEI). PEI (2.8 μg/well) was added dropwise to DNA (1 μg/well) in 150 mM NaCl and vortexed before adding the PEI/DNA mixture dropwise to the cell culture media. Cells were incubated 4 h at 37°C, 5% CO2, washed with 300 μl of fresh DMEM (including 10% HI-FBS) and maintained for a further two days. On the fourth day, cells were treated with recombinant human NKG2D-Fc chimeric receptor or Fc-control (1 μg/ml) in DMEM including 1% BSA, for 1h at 37°C, 5% CO2. Cells were then washed 3x PBS and fixed with PFA (4% diluted in 0.01M PBS) for 20-30 min before immunolabelling for human IgG (see NKG2D immunolabelling above).

### Confocal imaging

Immuno-labelled cultures were imaged on a laser scanning confocal microscope (LSM700, Zeiss) fitted with 3 laser lines (405, 488, and 546 nm). Individual neurons were first identified in the 405 nm laser channels corresponding to B-tubulin immunolabelling. Z-stack images (3-4 x 1 µm) of 405 nm, 488 nm and 546 nm channels were acquired at 1024 × 1024 resolution (12 bit) with a x20 air objective. 488 nm (Fc receptor) acquisition settings (i.e. gain, laser power, pin hole size) remained constant throughout all experiments.

For high-throughput quantification of NKG2D immunolabelling in sensory neuron cultures over time, coverslips were imaged on a spinning disc confocal microscope (iXplore SpinSR10, Olympus) with 4 Laser lines (405, 488, 561 and 640nm) fitted to an inverted microscope (IX83, Olympus) using stage navigator with Z-Drift Compensation for automated, systematic sampling. For mouse DRG neurons 100 regions of interest (ROI) were systematically sampled in a 10×10 grid aligned with the centre of the coverslip. For hiPSCd sensory neurons, 10 ROI were systematically sampled along the axis of laser ablation (see **Supp. Figure 6**). Z-stack images (3 x 2 µm) of 405 nm, 488 nm and 546 nm channels were acquired at 1156 x 1156 resolution (16 bit) with a x40 air objective. Identical laser and acquisition settings were maintained throughout.

An imaging pattern was designed for evenly taking ten images of both proximal and distal axons automatically by Spinning Disc confocal microscopy, which was applied to all the microfluidics. The distance between each ROI of proximal axions and microfluidics channel as well as between each ROI of proximal axons and the horizontally parallel ROI of distal axons was same. Each image containing 5 Z-slices with 2 µm step-size per slice was taken with a 40X objective. Zero-Drift Compensation (ZDC) was applied for maintaining the distance between objective and each ROI.

Super-resolution images of recombinant NKG2D receptor protein bound to *Mrgprd*+ DRG neurites were acquired on a Yokogawa CSU-W1SoRa Spinning Disc confocal microscope fitted with 4 Laser lines (405, 488, 561 and 640nm). 3D rendering and animation was performed using Zen 2012 software (Zeiss).

### Confocal image analysis

Analysis of NKG2D labelling of DRG neurons was performed using manual and automated methods. For manual counting, the number of NKG2D-binding neurons in the different subpopulations, individual β-tubIII+ neurons in a given field of view were manually assigned as either NKG2D-positive or negative while blinded to the expression of TdTomato. Images where neuronal densities were too high for identification of individual neurons (as assessed by the observer) were excluded from analysis.

For automated quantification of NKG2D labelling of murine DRG neurites, a bespoke analysis pipeline was executed using Fiji [48], in combination with a method for soma detection [49] and run via a script in MatLab with minor changes from the original publication to allow for batch processing of multiple images. The full script, including instructions for its use, can be downloaded from GitHub [50]. In brief, raw image files were imported to Fiji (ImageJ 2) by running Macro A, splitting into channels representing BtubIII (*blue*), NKG2D/Fc (*green*) and TdTomato (*red*) and generating pictures with a thresholded mask of the BtubIII channel representing total neuronal area. The newly generated image with a single channel BtubIII was imported to Matlab, where a soma mask array was generated via the Dimensionality Ratio method using the SomaExtraction package [49]. The mask array was imported back to Fiji to generate soma mask files by running Macro B. The soma mask was used to segment DRG neuron soma from neurites. Area selections (Regions of Interest, ROI) for βtubIII+ and TdTomato+ neurites were enlarged by 0.5 and transferred to the pre-thresholded NKG2D/Fc (*green*) channel for receptor quantification using the Particle Analysis function (size=0.1-400 µm^2^; circularity=0-1.00). Receptor density (particles per µm^2^) was calculated according to the original neurite area ROI for both βtubIII+ and TdTomato+ channels. A cut off of 200 µm^2^ was set for the minimum neurite area per ROI for receptor particle quantification. In experiments involving Thy1-YFP mice, the NKG2D and tdTomato fluorescence channels were reversed for quantitative analysis.

For automated quantification of NKG2D/Fc binding to human iPSC-derived sensory neurons, maximum intensity projection (MIP) images of BtubIII+ channel and the respective segmented images were generated by running Macro A. Debris with large size was identified by running SomaExtraction package on Metlab and Macro B on Fiji. The generated soma (debris) mask images were used for removing the noise from unspecific binding of NKG2D/Fc to debris generated by laser ablation. Area selections (Regions of Interest, ROI) for βtubIII+ were enlarged by 0.5 and transferred to the pre-thresholded NKG2D/Fc (*green*) channel for receptor quantification using the Particle Analysis function (size=0.1-400 µm^2^; circularity=0-1.00) by running Macro C. Receptor density (particles per µm2) was calculated according to the original neurite area ROI for BtubIII+ channel. The analysis pipeline can be found on GitHub [51].

The fragmentation of human iPSC-sensory neuron was quantified using semi-automated method [44]. The full Macro script can be found on GitHub [52]. Briefly, the area of axons (Area 1) and the area of BtubIII-positive particles (Area 2) in a same image were automatically measured by Macro script in Fiji. The fragmentation (%) = Area 1/Area 2 × 100.

### Human natural killer (NK) cell isolation and stimulation

Human NK cells were isolated from the peripheral blood of three healthy volunteers, as well as leukocyte cones from three volunteer blood donors supplied by UK National Health Service Blood and Transplant (NHSBT) Non-Clinical Issue (NCI) service.

Peripheral blood was collected by venepuncture in sodium-heparin tubes. Whole blood was then diluted 1:1 in serum-free RPMI 1640 media (Cat. 21875034, Life Technologies) and suspended on 15 ml Lympholyte Human Cell Separation Media (Cat. CL5020, CedarLane) in 50 ml SepMate tubes (Cat. 85450, Stem Cell Technologies) and centrifuged 22 min at 800g at room temperature (RT) with no brake to separate peripheral blood mononuclear cells (PBMC). Plasma containing platelets was removed and the leukocyte layer transferred to fresh RPMI containing 10% foetal bovine serum (heat-inactivated) (Cat. F9665-500ML, Sigma-Merck) and washed by pellet centrifugation at 500g, 10 min, RT. Live PBMC were counted by trypan blue exclusion and 10^8^ cells per sample were transferred to magnetic cell sorting (MACS) buffer (0.01M phosphate buffered saline (PBS) plus 2mM EDTA and 2% FBS). NK cells were enriched by MACS negative selection using a human NK Cell Isolation Kit (Cat. 130-092-657, Miltenyi Biotech) with LD columns (Cat. 130-042-901, Miltenyi-Biotech), according to the manufacturer’s instructions. Unlabelled NK cells were eluted into MACS buffer and counted with trypan blue exclusion for downstream applications. 4.65 – 6.75 × 10^6^ live NK cells were isolated per 10^8^ PBMC per donor. Approximately 10^6^ purified NK cells were sampled from each donor for purity check by flow cytometry analysis. Remaining NK cells were suspended at 10^6^ cells/ml in cryopreservation media (50% RPMI, 40% FBS, 10% DMSO) and frozen using a controlled-rate alcohol-free cell freezing container (CoolCell, Corning) before transfer to vapour phase nitrogen for long-term cryostorage.

Cone blood was diluted 1:1 in PBS containing 1% BSA and suspended on 15ml of Lympholyte Human Cell Separation Media (Cat. CL5020, CedarLane) in 50 ml Falcon tube and centrifuged 22 min at 800g (slow acceleration [3] and deceleration [0]) at room temperature (RT) to separate peripheral blood mononuclear cells (PBMCs). PBMCs located in the interphase (‘buffy coat’) between Lympholyte and plasma layers were carefully collected into a new 50ml Falcon tube and then washed by MACS buffer by pellet centrifugation 10 min at 400g at room temperature. 100×10^6^ PBMCs were resuspended in 2ml of MACS buffer. NK cells were enriched by negative selection using EasySep^TM^ Human NK Cell Isolation Kit (Stem Cell, Catalog #17955), according to the manufacturer’s instructions. The isolated NK cells were suspended in cryopreservation media (50% RPMI, 40% FBS, 10% DMSO) and frozen using a controlled-rate alcohol-free cell freezing container (CoolCell, Corning) in −80°C before being transferred to vapour phase nitrogen for long-term cryostorage.

For NK cell stimulation, vials of cryopreserved NK cells were rapidly thawed in a water bath at 37°C and washed in RPMI including 10% FBS supplemented with DNase I (125 U/ml) followed by centrifugation at 400g, 10 min, RT. Cells were counted and seeded at 2×10^6^ cells per ml in RPMI+10% FBS in 96 well U-bottom plates (Nunclon Delta, Cat. 10344311, FisherScientific) supplemented with recombinant human IL-2 (10^3^ U/ml; 100ng/ml equivalent) (Cat. 200-02, Peprotech) and cultured for 2 days at 37°C, 5% CO_2_.

### Flow cytometry

Whole PBMC, NK-depleted fraction and purified human NK cells (5×10^5^ cells per 100 µl) were suspended in FACS buffer (PBS +2% FBS) and blocked with normal human serum (NHS, 10%) for 15 min on ice. Cells were treated with fluorescently-conjugated antibodies (see **Supp. Table 4**) and incubated 40 min at 4°C protected from light. Cells were washed 2x FACS buffer with centrifugation 500g, 5 min. The fluorescent DNA intercalator 7-aminoactinomycin D (7-AAD) (1:100) was added to all samples (except single stain controls) prior to cytometry. Samples underwent flow cytometry on an LSRII Special Order Research Product (SORP) digital cell analyser equipped with a Violet (405nm, 100mW), Blue (adjustable 488nm, 80mW), Green (532nm, 150mW) and Red (642nm, 40mW) lasers. Prior to data acquisition the cytometer was calibrated using CS&T Beads. PMT gain and compensation settings were established using single and unstained control PBMC samples. Due to limited spill-over between fluorophores, fluorescence-minus-one controls were not employed. Approximately 100,000 events were run per sample. Data were exported as .fcs files and analysed in FlowJo. Lymphocytes were gated based on characteristic forward and side scatter. Marker gates were set based on single cell controls. The gating strategy was as follows: Lymphocytes -> 7AAD^neg^ -> singlets -> CD19^neg^CD3^neg^ -> CD56^dim^CD16^+^ (cytotoxic NK) versus CD56^bright^CD16^neg^ (regulatory NK). NKp46 and NKG2D receptor expression levels, as well as production of perforin and granzyme B by NK cells were analysed by full spectrum flow cytometry (Aurora 5 laser Spectral Flow Cytometer, Cytek) equipped with UV (355nm, 20mW), violet (405nm, 100mW), blue (488nm, 50mW), 50mW yellow-green & red (638nm, 80mW) lasers. Quality control (QC) was performed by running SpectroFlo QC beads before acquiring samples, automatically optimizing laser and the gains (voltages) for every detector. Then FSC (Forward Scatter) and SSC (Side Scatter) were adjusted to ensure that cells were not off scale. Reference controls without staining or single staining only were acquired for spectral unmixing and FMO (fluorescence minus one) controls were prepared for defining the boundaries of positive and negative cell populations. Raw data were unmixed by SpectroFlo and the unmixed data were analysed by FlowJo (version: 10.10.0). 1×10^6^ cells were resuspended in 100 µl FACS buffer (PBS containing 2% FBS) were incubated with LIVE/DEAD Fixable Blue (Cat. L34961, Biolegend) 20 min on ice protected from light. Cells were washed twice with FACS buffer. Cells were incubated with 10% normal mouse serum on ice for 15 min and then incubated with anti-CD45 (Cat. 563792, BD Biosciences, RRID: AB_2869519), anti-CD3 (Cat. 612941, BD Biosciences, RRID: AB_2916883), anti-CD14 (Cat. 301831, Biolegend, RRID: AB_2563629), anti-CD11c (Cat. 612967, BD Biosciences, RRID: AB_2870241), anti-CD56 (Cat. 318333, Biolegend, RRID: AB_2561912), anti-CD16 (Cat. 741449, BD Biosciences, RRID: AB_2870923), anti-NKG2D (Cat. 320818, Biolegend, RRID: AB_2562792) for 45 min on ice protected from light (see **Supp. Table 4**). Cells were washed twice with FACS buffer and then fixed with Fixation and Permeabilization Solution (Cat. 554714, BD Biosciences) on ice for 20 min. Cells were washed twice with Perm/Wash Buffer (BD Biosciences) and incubated with anti-perforin (Cat. 48-9994-42, Life Technologies, RRID: AB_2574145) and anti-granzyme B (Cat. 58-8896-42, Life Technologies, RRID: AB_2724390). Cells were washed twice with Perm/Wash Buffer (BD Biosciences) and resuspended in 100 µl FACS buffer for full spectrum flow cytometer analysis.

### NKG2D blocking assay

For antibody blockade of human NKG2D function *in vitro*, NK cells were incubated with 50 μg/ml of LEAF purified anti-human CD314 (NKG2D) (clone 1D11) (Cat. 320813, Biolegend; RRID: AB_2561488) or LEAF purified human IgG1 isotype control (Clone QA16A12) (Cat. 403501, Biolegend; RRID: AB_2927629) for 30 min at room temperature (2.5×10^6 NK cells per ml) before addition to target cells (hiPSCd sensory neurons) in neurite compartment of microfluidic devices. Cells were seeded at different densities (50×10^3^, 100×10^3^, 200×10^3^) for 4 h.

NK cells were added to the neurite compartment of microfluidic devices approximately 4 weeks after hiPSCd-sensory neuron seeding and co-cultured for 4 h. Then cells were washed three times by HBSS, followed by fixation by 4% PFA for 30 min. Fixed cells were washed three times by HBSS and blocked and permeabilized by PBS containing 5% normal serum and 0.1% Triton-X100 for 1 h at room temperature. Then cells were incubated with primary antibody diluted in PBS containing 0.5% normal serum and 0.01% Triton-X100 overnight at 4°C. Cells were washed three times by PBS and incubated with DAPI and secondary antibody diluted in PBS containing 0.5% normal serum and 0.01% Triton-X100 for 1 h at room temperature. Then cells were washed three time by PBS and the reservoirs of microfluidics were filled up with PBS and covered with coverslip.

### Quantitative PCR

#### RNA extraction from mouse DRG cultures

After various time points in culture (day 1, 2, and 3), coverslips were gently washed with PBS before lysis with a phenol-containing lysis buffer (Tripure, Roche). Lysates were transferred to 1.5 mL samples tubes, snap-frozen on dry ice, and stored at −80°C until RNA extraction using filter column purification (High Pure RNA Tissue Kit, Cat. 12033674001, Roche).

Samples from the same batch of cultures were processed simultaneously by thawing at room temperature (RT) for 5 min. 100 µL of chloroform was added to each sample, samples were shaken for 15 s, then incubated at room temperature for 15 min. Samples were centrifuged at 12,000G for 15 min at 4°C. 200 µL of the upper aqueous phase was added to 200 µL of 70% ethanol in a filter column, which was centrifuged at 13,000G for 30 s. DNase solution was added, and the column incubated at RT for 15 min. RNA was purified by a series of wash steps according to the manufacturer’s instructions followed by elution buffer and centrifugation at 8,000G for 1 min. RNA eluates were checked for concentration and purity on a NanoDrop Microvolume Spectrophotometer and stored at −80°C until reverse transcription.

#### Reverse transcription

Reverse transcription for quantitative PCR was performed using Moloney Murine Leukemia Virus (M-MLV) reverse transcriptase kit (Cat. 28025-013) with oligo(dT) primers (Cat. 58862) (Invitrogen, Thermofisher). Briefly, 50 ng RNA was added to 1 µL Oligo(dT)12-18 (500 µg/mL), and 1 µL 10mM dNTP in PCR-grade water in a nuclease-free reaction tube (total volume 12 µL). The mixture was heated to 65°C for 5 min then quick-chilled on ice. To each tube was added: 4µL 5X First-Strand Buffer, 2 µL 100mM DTT, and 1 µL RNase OUT (40 units/ml) (Cat. 100000840) and mixed gently. Samples were incubated at 37°C for 2 min before adding 1 µL (200 units) of M-MLV RT to each tube, mixing by gentle pipetting and incubation at 37°C for 50 minutes, before heating at 70°C for 15 min. Complimentary DNA (cDNA) products were stored at −20°C until use in PCR experiments. Universal Mouse Reference RNA (Cat. QS0640, Life Technologies) and DRG-extracted RNA with omission of reverse transcriptase were used as positive and negative controls, respectively.

#### Quantitative polymerase chain reaction (qPCR)

Quantitative gene expression in DRG cell cultures was performed on cDNA (0.33 ul per tube per sample from original reverse transcription) using a SYBR Green PCR Master Mix (Roche) and pairs of target-specific primers (500 nM each) in a 10 µl sample volume on white skirted 384-well PCR plate (Cat. E1042-9909, StarLab) on a LightCycler 480 (Roche). The PCR conditions were: 95°C (5 min), and cycled 45 times at 95°C (30 s) to 60°C (2 min) followed by melt curve for PCR product confirmation: 65°C (1 min), ramped to 97°C (0.11 °C/s). Gene expression was analysis performed on cDNA from DRG using a Power SYBR Green PCR Master Mix (Applied Biosystems) and pairs of target-specific primers (500nM) in MicroAmp optical tubes (20µl reaction volume) on a 7500 Real-Time PCR system (Applied Biosystems). The PCR conditions were 50°C (2 min), 95°C (10 min), and cycled 40 times at 95°C (15 s) to 60°C (1 min). Data were analysed using the built-in 7500 software (v2.0.4, Life Technologies). All samples were run in triplicate. Expression was determined relative to a reference gene (Gapdh) using the comparative Ct method [53]: 2^-ΔΔCt^ = 2^-(Ct of Gapdh – Ct of target gene^). ΔΔCt values were normalised to day 0 DRG to present as fold change in transcripts at each time point for each culture, or to contralateral DRG for nerve injury experiments.

Primers were designed using PrimerBLAST [54] or obtained from previous literature (See **Supp. Table 5**). Primers had GC content between 40% and 60%. Where possible, primers were designed to overlap the exon-exon boundary, and 3’ ends were G or C Annealing temperatures of primer pairs were within 5°C of each other. OligoCalc [55] was used to select primers with minimal potential for hairpin formation, 3’ complementarity and self-annealing, and were validated by a single peak in the dissociation curve or single band PCR product via gel electrophoresis.

#### Gel electrophoresis

qPCR reaction products (10 µL) were run on 1.5% agarose gel stained with GelRed (Cat. SCT123, Merck Millipore) at 100V for 30 min with 100 bp ladder (Cat. N3231S, New England Biolabs). The gels were imaged using a UV illuminator (NuGenius, Syngene).

### Single cell DRG collection, reverse transcription and nested PCR

Adult male C57BL/6 mouse DRG neurons were cultured as above overnight in Neurobasal media supplemented with NGF (50 ng/ml) on coverslips previously coated with poly-D-lysine (10µg/ml) and laminin (10µg/ml). Coverslips were transferred to a patch-clamp recording chamber and perfused with a 2Ca^2+^/Na^+^ buffer containing (in mM): 140 NaCl, 10 HEPES, 2 CaCl_2_, 1 MgCl_2_, 10 glucose, 5 KCl, pH 7.2 in DEPC-treated water. Micropipettes for cell collection with a tip diameter of approximately 10 µm were prepared from borosilicate glass previously washed with DEPC-treated water and autoclaved to eradicate contaminating RNase activity. Single DRG neurons were collected into the pipette containing a reverse transcription buffer by applying gentle negative pressure under visual control and were immediately ejected into a collection buffer containing dNTP, oligo(dT) and random hexamers. Negative controls were provided by collection of the perfusion solution only. Samples were heated to 65°C for 5 min and cooled on ice before the addition of reaction buffer containing reverse-transcriptase (Superscript III, Invitrogen, Life Technologies). cDNA was synthesised on a thermo-cycler (Biorad T100) (25°C, 5 min; 50°C, 100 min; 85°C, 5 min) and finally 37°C for 20 min in the presence of RNase H. cDNA was stored at −20°C. PCR was performed with nested primers using Platinum *Taq* DNA polymerase (Invitrogen) according to the manufacturer’s instructions. The first round of PCR was performed with 1-1.5 µl single cell cDNA and ‘outer’ primer pairs (95°C, 5 min followed by melting at 94°C for 40s, annealing at 52-60°C for 40s and elongation at 72°C for 40s, cycled 35 times). The second round of PCR was performed with 2 µl of the first round PCR product, ‘inner’ primer pairs and cycled 20 times. Second round PCR products were visualised by gel electrophoresis stained with SafePinky (GenDepot). Nested primer pairs were synthesised by a commercial supplier (Bioneer, Korea) (see **Supp. Table 6**).

### RNA sequencing dataset analysis

Human DRG (hDRG) sequencing data were downloaded as transcript-per-million (TPM) counts for Wangzhou et al (2020) [56] and quantile normalized TPM (qnTPM) from Ray et al., 2023 (supplemental table 2, quantile normalized sheet) [57]. In line with published their published reports, only hDRG with clear neuronal enrichment were used from Ray et al., 2023. Ensembl IDs were mapped to gene symbols using biomaRt in R.

Visium data were accessed as a Seurat Object from Tavares-Ferreira et al. (2022) [58]. Single soma hDRG count data were downloaded from GEO (GSE249746) [59]. iPSC (AD2, AD3 and 840 lines) and iPSC-derived sensory neuron data were downloaded from GEO (GSE144208) [60].

Non-neuronal cells from a cross-species harmonized DRG atlas were downloaded from the ‘painseq’ dataset [61] as a Seurat object. Human barcodes were extracted, and the object was subset for the “Immune” cluster. The extracted RNA counts were used to create a fresh object prior to downstream processing in Seurat. Data were normalized (‘NormalizeData ‘), and variable features extracted ‘FindVariableFeatures(seu.immune, selection.method = “vst”, nfeatures = 2000)’. After scaling, and elbow plot was used to determine dimensions (here, 10). Clustering was performed as ‘FindClusters(seu.immune, resolution = 1)’. The original published object was mapped to mouse gene names, thus this naming was maintained for plotting.

Human sural nerve data was accessed from the ‘osmzhlab’ dataset [62]. Mouse DRG neuron data was accessed from the ‘ernforslab’ dataset [63]. Plots were downloaded as png files from the corresponding Shiny application.

Data were processed in line with their publication. Briefly, Seurat objects were created for individual donors (n=6) in R, and normalized (‘NormalizeData(seurat_obj)’. Variable features were calculated on the top 4500 features (‘FindVariableFeatures(seurat_obj, nfeatures = 4500)’). Integration anchors were calculated using FindIntegrationAnchors(object.list = seurat_objects, scale=TRUE, normalization.method=“LogNormalize”, reduction=“cca”, l2.norm = TRUE, dims=1:30, k.anchor=5, k.filter=200, k.score=30, max.features=200, nn.method=“annoy”, n.trees=50, eps=0). Objects were then integrated using IntegrateData(ss.data, normalization.method=“LogNormalize”, features=NULL, features.to.integrate=NULL, dims=1:30, k.weight=80, weight.reduction=NULL, sd.weight=1, sample.tree=NULL, preserve.order=FALSE, eps=0). Data were then scaled (‘ScaleData ‘) prior to dimensionality reduction (RunPCA(ss.data, npcs=50), RunUMAP(ss.data, reduction=“pca”, dims=1:25)). Clusters were then calculated using ‘FindNeighbors(ss.data, reduction=“pca”, dims=1:25)’ and ‘FindClusters(ss.data, resolution=3.4)’ in line with published methods from Yu et al. (2024). Metadata were extracted from the project-associated github (taimeimiaole/NN_hDRG-neuron-sequencing/Source_code_2/human_meta_final_cluster.Rdata), with ‘cl.conserv_final’ set as Idents for plotting.

All plots were generated in R, using ggplot2 and/or Seurat.

### Study design and statistics

For *in vivo* experiments, biological unit of interest is the animal (i.e. number of mice). For murine cell culture experiments the biological unit of interest is either the animal (i.e. number of cultures for RNA analysis) or the cell (i.e. number of DRG neurons for immunohistochemical analysis), unless otherwise defined; where individual neurons were not defined (i.e. during automated image analysis), the biological unit was a region of interest. Mice of both sexes were used unless otherwise indicated. For human cell culture experiments, the biological unit of interest is the individual donor. Experimental units are defined in figure legends. The group containing devices without NK cells were excluded from the data analysis because of the low density of axons. Where multiple observations were made of RNA levels or immunohistochemical signals *in vitro*, the various treatments (including controls) were applied to replicate experimental units (i.e. cell culture well) derived from each biological unit.

Each replicate cell culture contained the full suite of experimental and control samples. Assignment of cell culture wells to different time points or treatment with different reagents (i.e. recombinant proteins or cells) was not randomised. Counting of neurons was performed on images offline after manual image acquisition. DRG neuron culture from a single male mouse for resulted in low levels of extracted RNA (<5 µg/ml) at days 1, 2 and 3 *in vitro* **Fig. 2B**), therefore data from this animal was excluded from analysis.

For *in vivo* experiments, sample sizes (i.e. number of animals) were based on previous publication [18]. *In vitro* experiments were exploratory and therefore sample size was not calculated *a prior*. All the graphs, calculations and statistical analyses were performed using GraphPad Prism software 10.0. Data points represent mean values of replicate measurements with standard error of the mean, unless otherwise stated. Pair-wise comparisons of normally distributed data were analysed using a two-tailed Student’s t test. Pair-wise comparisons of non-normally distributed data were analysed using a two-tailed Mann-Whitney U test. Where the experimental unit is a single image (i.e. when calculating receptor particle density) data were compared as cumulative distributions using the Kolmogorov-Smirnov test and presented as violin plots showing median and quartile ranges. Alpha=0.05.

All data points were included in the data analysis except for the excluded data points described above.

## RESULTS

### DRG neuron culture replicates injury-induced regulation of NKG2D ligands

We first sought to identify the *Raet1* gene isoforms expressed by DRG neurons. Transcripts for *Raet1*d and *Raet1*e [27,64] were detected in DRG tissue, while transcripts for *Raet1a*, *Raet1b* and *Raet1c* were not detected **Fig. 1A**), consistent with previous findings in other tissues from C57BL/6J mice [33,65]. Quantitative PCR of DRG cultures revealed an upregulation of *Raet1*e, and to a lesser extent, *Raet1*d, over several days in vitro, consistent with primers against all *Raet1* isoforms (**Fig. 1B**) [18]. DRG neuron culture also led to an upregulation of the nerve injury-related transcription factor *Atf3* (**Fig. 1B**) suggesting that DRG neurons *in vitro* represent an injury-like state [12]. Cell culture had no effect on the ratio of house-keeping genes *Gapdh* and *Actin* (**Fig. 1B**, *right*)

**Figure 1.**
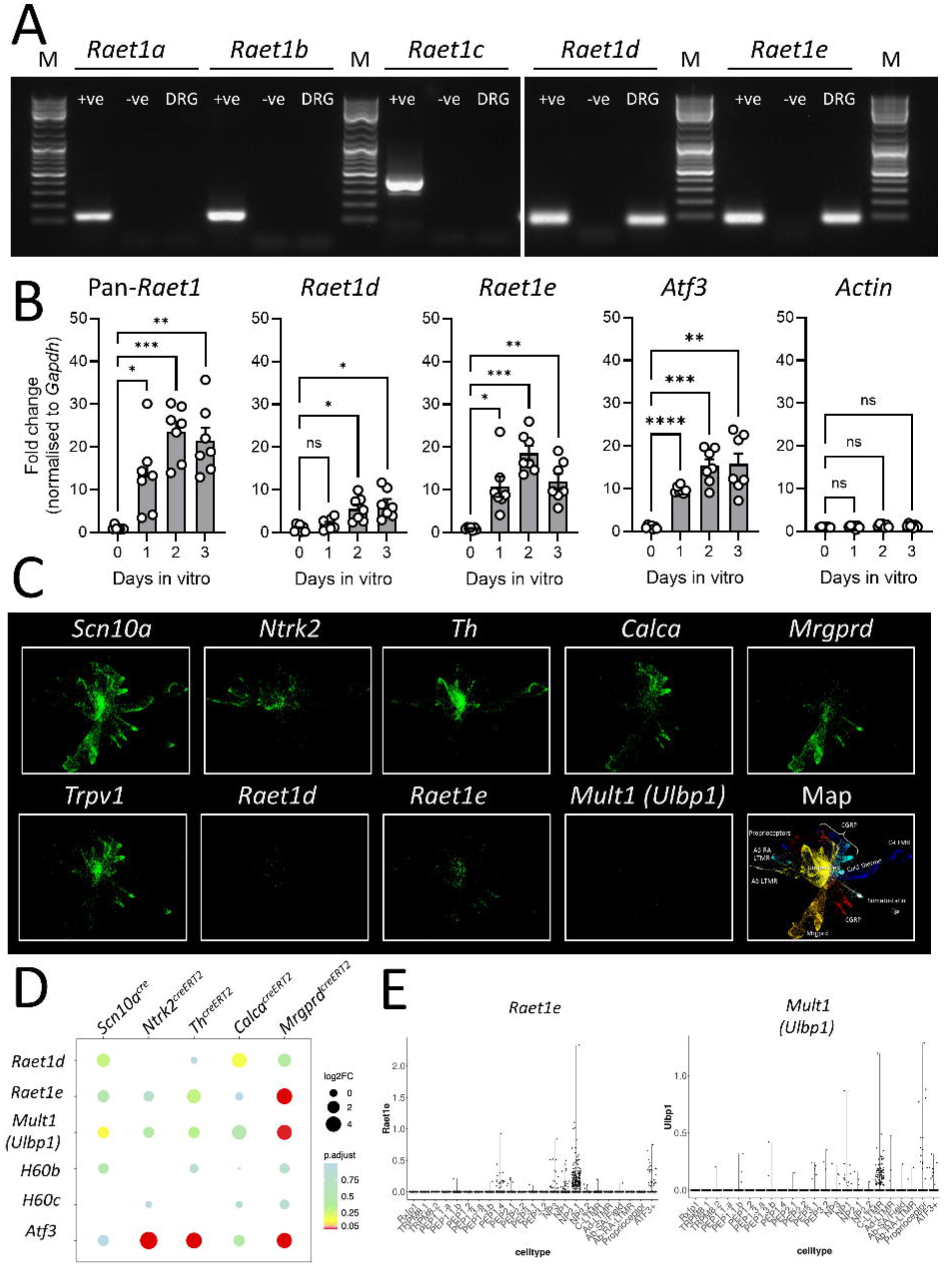
Detection of mRNA transcripts for Raet1 gene family in murine DRG neuron. **A)** PCR gel electrophoresis showing detection of *Raet1δ* and *Raet1ε* mRNA transcripts only in DRG from C57BL/6J mice. Predicted PCR product sizes for *Raet1α* (111 bp), *Raet1β* (119 bp), *Raet1γ* (404 bp), *Raet1δ* (67 bp), *Raet1ε* (79 bp). +ve, universal RNA positive control; -ve, minus RT control reaction; DRG, lysate from DRG neurons after 3 day in vitro (female mouse); M, 100 bp DNA ladder marker. **B)** qPCR for pan-*Raet1, Raet1δ*, and *Raet1ε* mRNA transcripts in DRG cell cultures over 3 days in vitro. *Gapdh* was used a reference gene. n=7 replicate DRG cultures from different mice (3m, 4f). Repeat measures ANOVA versus Day 0: *pan-Raet1,* F_(2.547, 15.28)_ = 19.67, P<0.0001; *Raet1δ,* F_(1.864, 11.19)_ =10.51, P=0.0030; *Raet1ε*, F_(1.696, 10.17)_ = 24.34, P=0.0002; *Atf3*, F_(1.079, 6.472)_ = 31.19, P=0.0010; *Actin*, F (2.361, 14.17) = 2.881, P=0.0825. Dunnett’s multiple comparison (corrected p values): *p<0.05, **p<0.01, ***p<0.001. **C)** Visualization of force-directed layout of mouse DRG sensory neurons overlaid with single cell RNA expression of lineage marker genes *Scn10a (*general nociceptors), *Calca* (peptidergic nociceptors), *Mrgprd* (non-peptidergic nociceptors), *Th* (C-LTMR) and *Ntrk2* (Aβ-RA + Aδ-LTMR), Trvp1 (Thermosensitive nociceptors) and NKG2D ligands *Raet1d*, *Raet1e* and *Mult1* (*Ulbp1*). Data from Sharma et al. 2020 [67]. **D)** NKG2D ligand expression in each of five sensory neuron lineages after nerve injury. Gene expression levels are presented as transformed transcript counts in ipsilateral and contralateral lumbar DRG neurons 3 days after spared nerve injury. Data from Barry et al., 2023 [41]. **E)** Violin plots showing normalised expression of *Raet1e* and *Mult1* (*Ulbp1*) among mouse DRG neuronal subtypes. Data from integrated atlas curated by Krauter et al. [70].

*H60a* is not detected in C57BL/6 mice (Lodoen et al., 2003) and with the exception of the skin, transcripts for H60b and H60c are relatively low [66]. We therefore investigated the expression of the high affinity *Mult1* ligand after peripheral nerve injury. Interestingly, we observed an increase in *Mult1* expression in ipsilateral L5 DRG after spinal nerve transection (**Supp. Figure 1A**), similar to previous findings with *Raet1* [18]. In contrast, expression of the inhibitory MHC class I molecule *Qa1b* was not significantly regulated in DRG by nerve injury, while the MHC class I adaptor molecule β2-microglobulin (*B2m*) showed a small but significant increase in DRG 7 days after L5x compared to sham (**Supp. Figure 1B**).

To delve deeper into the sensory neuron subsets expressing NKG2D ligand genes we explored published single cell datasets. Consistent with our PCR results in C57BL/6 mice, transcripts for *Raet1d*, *Raet1e* and *Mult1* (*Ulbp1*), but not *Raet1a*, *Raet1b* and *Raet1c,* were among those detectable in DRG neurons from naïve mice [67–69]. As expected, RNA levels in healthy tissues were low, precluding a quantitative assessment of enrichment in one or other sensory neuron subtype as defined by canonical marker genes (**Fig. 1C**). To overcome the possible limitations in sequencing depth of single cell datasets, we analysed a pseudo-bulk RNA sequencing dataset comparing gene expression from pooled single DRG neurons 3 days after peripheral nerve injury [41]. Among the known NKG2D ligands detectable in the dataset of five different labelled sensory neurons lines, *Raet1e* and *Mult1* (*Ulbp1*) transcripts were most abundant and significantly expressed in *Mrgprd*-expressing neurons ipsilateral to nerve injury (**Fig. 1D**). A recently published integrated atlas of over 44,000 mouse DRG neurons from several datasets [70] showed *Raet1e* clustering within non-peptidergic DRG neurons, while Ulbp1 (*Mult1)* was enriched predominantly in *Th*+ C-low threshold mechanoreceptors (C-LTMRs) (**Fig. 1E**). In contrast, *Raet1d, H60b* and *H60c* were detected at very low levels (**Supp. Figure 1C**), while *Raet1a*, *Raet1b*, *Raet1c* and H60a were undetected (data not shown), as expected due to the majority of published studies using mice on a C57BL/6 background.

### Detecting NKG2D ligand genes at single DRG neuron resolution

To validate the expression of NKG2D ligands at the single cell level, and to ask whether transcript prevalence was affected by prior nerve injury, we collected individual DRG neurons under microscopic control from acute cultures of DRG neurons (<24 h *in vitro*) from adult C57BL/6 mice 7 days after sham or L5-spinal nerve transection (L5x) surgery. Collected cells were lysed and underwent reverse-transcription followed by two rounds of nested PCR with gene-specific primers (See Methods). A total of 18 sham and 20 L5x cells out of 24 single cells collected per group were positive for *Gapdh* and *Advillin* transcripts indicating successful DRG soma collection (**Figs 2A, B** and **Supp. Figures 2A, B**). Due to a limited amount of PCR product, we limited our analysis to a small group of genes. Pan-*Raet1* transcripts were observed in 20-30% of identified neurons (**Fig. 2B** and **Supp. Figures 2A, B**), with no difference in frequency between cultures from sham and L5x injured mice (p>0.999, Fisher’s exact test). Only a small number of neurons (5-10%) co-expressed transcripts for both *Raet1* and *Trpv1*. In sham DRG cultures all *Raet1* and/or *Trpv1*-expressing neurons were positive for *Scn10*a (**Fig. 2C**, *left*), the gene encoding the voltage-gated Na^+^ channel Na 1.8. Fewer neurons from L5x DRG were positive for *Scn10*a (**Fig. 2C**, *right*) consistent with the down-regulation of Na_v_1.8 after nerve injury [13]. Detection of *Mult1* mRNA transcripts in DRG neurons at the single cell level - either sham or L5x injured - was rare in our sample population (**Fig. 2C** and **Supp. Figures 1A, B**).

**Figure 2.**
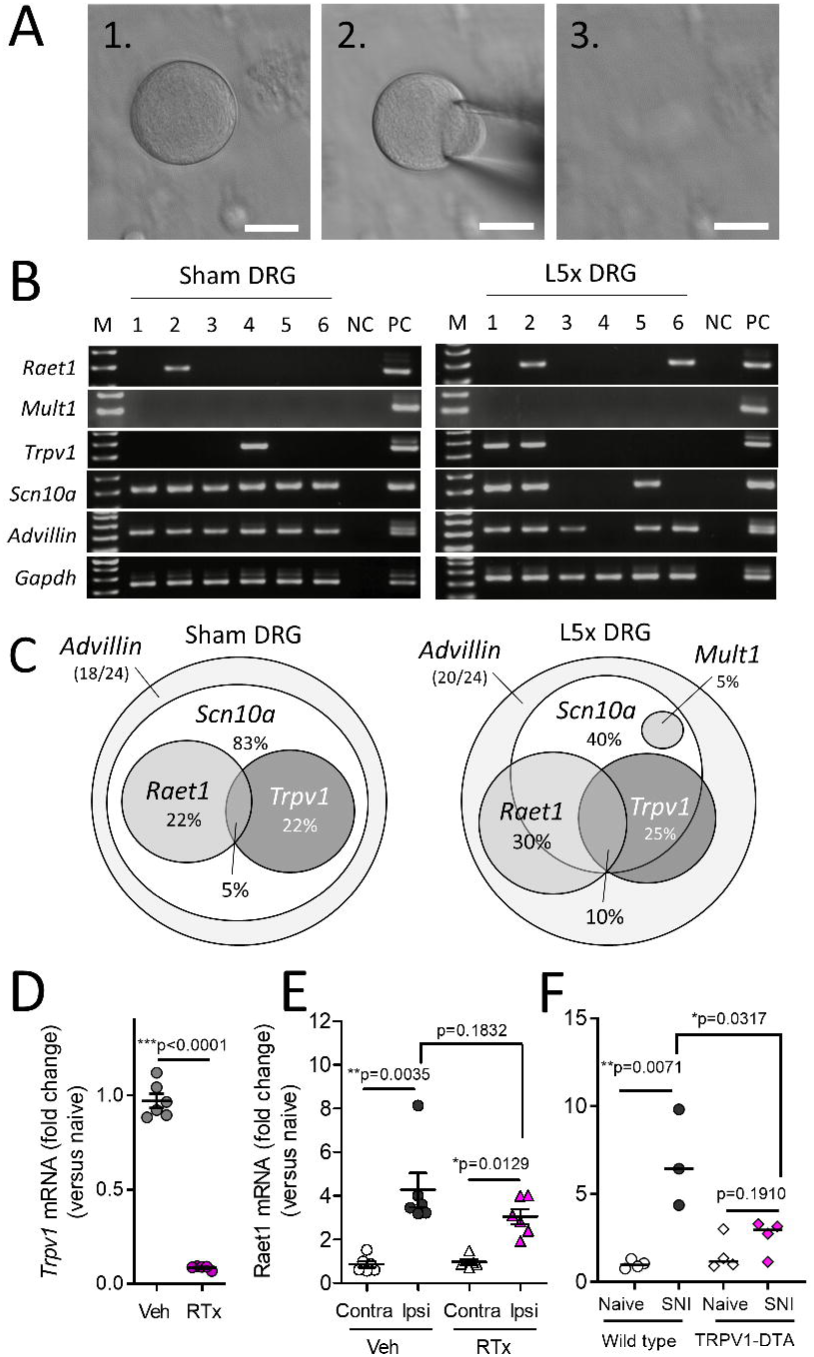
Raet1 is expressed by individual nociceptive DRG neurons. **A)** Consecutive images of single DRG neuron collection by glass pipette (scale bars, 10 µm). **B)** Single cell nested PCR results for multiple mRNA transcripts detected in DRG neurons isolated from adult male C57BL/6J mice 7 days sham (uninjured) (*left*) or lumbar L5 spinal nerve transection (L5x) surgery (*right*) <24h *in vitro.* PCR bands for six DRG neurons are shown with DNA ladder marker, M. NC, negative control (bath solution). PC, positive control (whole DRG tissue). **C)** Venn diagrams showing the proportion of *Gapdh+Advillin+* DRG neurons with transcripts detected for *Scn10a* (Nav1.8), *Raet1* and *Trpv1,* 7 days after sham surgery (*left*) or L5x-injury (*right*). Central overlapping region represents *Raet1*+*Trpv1+* double-positive DRG neurons. **D)** qPCR showing the effect of neonatal resiniferatoxin (RTx) treatment on *Trpv1* mRNA expression in lumbar L3-5 DRG compared to vehicle. Student’s unpaired t test: t=20.99, p<0.0001. (**E**) qPCR shows crush injury-related increase in *Raet1* mRNA expression in ipsilateral L3-5 DRG 7 days after surgery both in vehicle and RTx-treated adult male C57BL/6J mice. Student’s paired t test: Vehicle, contra *versus* ipsi, t=5.200, **p=0.0035; RTx, contra *versus* ipsi, t=4.278, *p=0.0129; Ipsi, Veh *versus* RTx, t=1.430, p=0.1832. n=6 mice per treatment. (**F**) Increase in *Raet1* mRNA expression in ipsilateral L4-5 DRG 7 days after spared nerve injury (SNI) is attenuated in *Trpv1^cre/wt^;rosa26^dta/wt^* mice. Student’s unpaired t test: SNI *versus* naïve wild type, t=4.395, p=0.0071; SNI *versus* naïve *Trpv1-dta*, t=1.474, p=0.1910. n=3-4 mice per surgery per genotype.

To confirm the identity of small-sized neurons in the nerve injury group we additionally checked for expression of the transcription factor *Runx1* and nerve growth factor (NGF) receptor *Trka* [71]. All *Raet1* and *Trpv1* neurons were positive for *Runx1,* while only around 50% were positive for Trka (**Supp. Figures 2C, D**). The proportion of *Runx1* and *Trka* transcripts among *Raet1*-expressing neurons was consistent between sham (**Supp. Figure 2C**) and L5x DRG (**Supp. Figure 2D**). The transcriptional profile of *Raet1* neurons (expressing *Runx1* and partial overlap with *Trka* and *Trpv1*) was in keeping with the characterisation of *Trpv1*-lineage neurons [72]. Chemical ablation of TRPV1-expressing cells by neonatal treatment with the potent agonist resiniferatoxin (RTx) (**Fig. 2D** and **Supp. Figure 2E**) reduced but did not completely prevent the nerve injury-induced expression of *Raet1* in ipsilateral DRG tissue (**Fig. 2D, E**). On the other hand, crossing a *Trpv1*-driven Cre-recombinase expressing mouse (*Trpv1^cre^*) [38] with a diphtheria toxin subunit A (DTA) reporter mouse to ablate the *Trpv1*-lineage neurons prevented the upregulation of *Raet1* in DRG 7 days after peripheral nerve injury (**Fig. 2F**), suggesting the majority of *Raet1*-expressing neurons lie within the *Trpv1*-lineage nociceptive population.

In summary, *Raet1e* and *Mult1* (*Ulbp1)* were the two main NKG2D ligand transcripts enriched in unmyelinated sensory neurons and upregulated by peripheral nerve injury. While *Raet1e* is predominantly restricted to a subpopulation of nociceptive DRG neurons characterised by the developmental expression of *Scn10a* and *Trpv1*, *Mult1* (*Ulbp1)* appears to have a more limited RNA expression, possibly in C-LTMRs.

### NKG2D ligand detection in a live cell-based assay

To demonstrate functional relevance of these findings, we next sought to validate the detection of NKG2D ligands at the protein level using a live cell-based assay approach. The NKG2D ligand family was originally discovered in mouse tumour cell lines with the use of NKG2D-Ig fusion proteins [27]. We first validated a source of soluble NKG2D receptor protein chimerised to the Fc domain of human IgG1 (NKG2D-Fc) to recognise the high affinity ligand mouse Raet1e by heterologous expression in HEK293T cells (**Supp. Figure 3A**). Incubation of live cells with 2 µg/ml of NKG2D-Fc chimeric protein, but not an equivalent amount of Fc-only control protein, bound to the cell surface of cells transfected with *Raet1e* and not untransfected controls (**Supp. Figure 3B**).

Application of NKG2D-Fc to live DRG neurons (3 days *in vitro*) under the same conditions, revealed punctate labelling of some, but not all, BtubIII+ neurites (**Fig. 3A**). NKG2D receptor binding to DRG neurites increased over time (**Fig. 3B**), paralleling the upregulation of *Raet1e* transcript (**Fig. 1B**). Quantification of immunofluorescence signal density of confirmed significant detection of NKG2D-Fc receptor binding versus Fc controls by 3 days *in vitro* (**Fig. 3C, D**).

**Figure 3.**
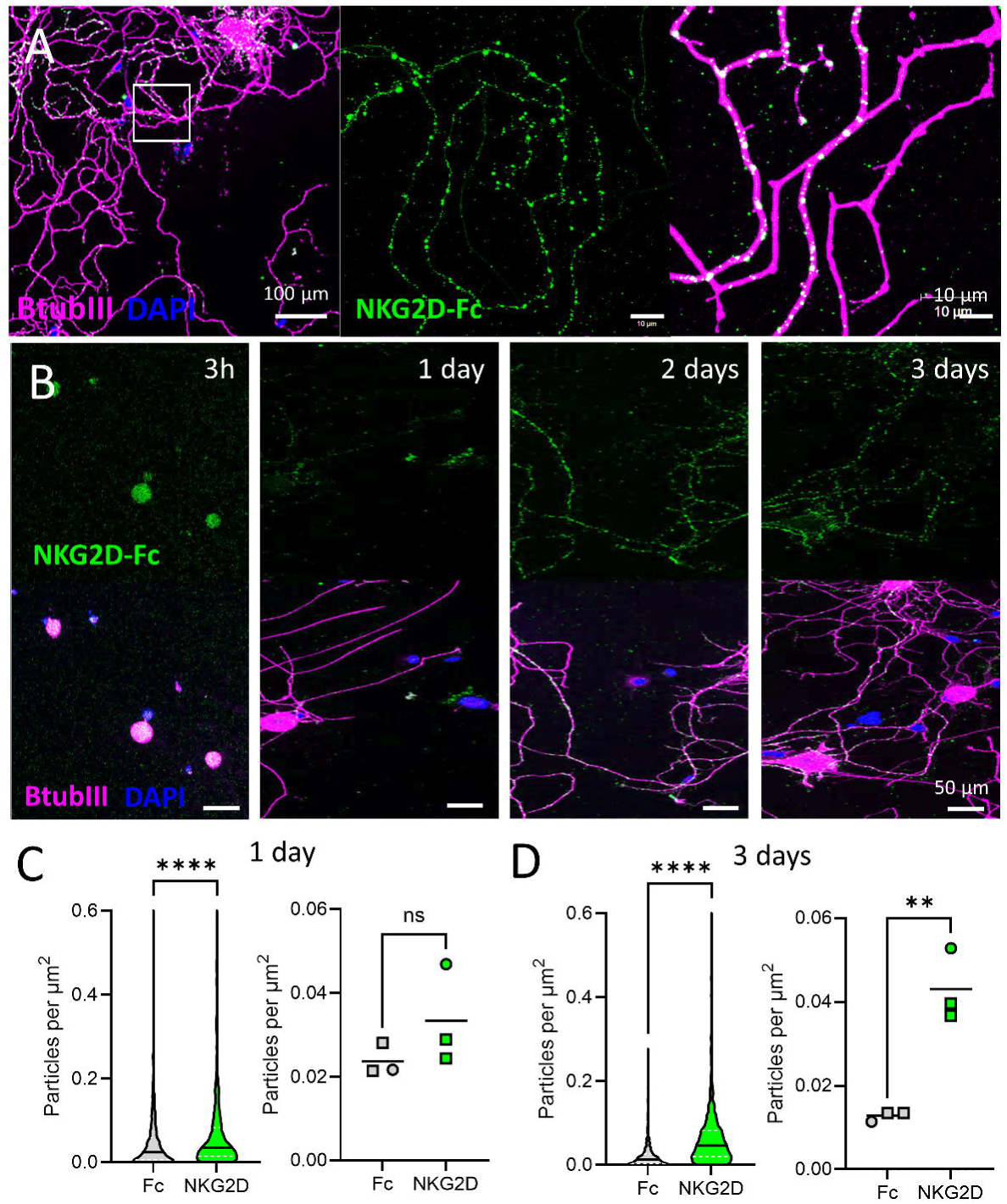
Soluble NKG2D receptor binding to DRG neurite membrane increases over time in culture. **A)** *Left*) Low magnification view of DRG neurons 3 days *in vitro*. *Right*) High magnification of inset NKG2D-Fc chimeric protein labelling (*green*) of BtubIII+ neurites (*magenta*). DAPI, *blue*. **B)** Representative images of NKG2D-Fc binding to DRG neurons over time in culture. Scale bars, 50 µm. **C)** Quantification of NKG2D-Fc binding versus Fc-only control on day 1 *in vitro*. *Left*) Violin plot of particle density per µm^2^ of neurite area per image: Fc, n=609 images; NKG2D-Fc, n=661 images. ****p=<0.0001, Kolmogorov-Smirnov test (D=0.1389). *Right*) Median particle density per mouse; ns, p=0.2512, t=1.340, two-tailed unpaired t-test. **D)** Quantification of NKG2D-Fc binding versus Fc-only control on day 3 *in vitro. Left*) Violin plot of particle density per µm^2^ of neurite area per image: Fc, n=707 images; NKG2D-Fc, n=769 images. ****p=<0.0001, Kolmogorov-Smirnov test (D=0.4296). *Right*) Median particle density per mouse; **p=0.0038 (t=6.042) two-tailed unpaired t-test. n=1f (*circle*), n=2m (*squares*) mice per time point.

### NKG2D receptor binding to sensory neuron subsets

We next asked whether NKG2D-Fc receptor binding preferentially targeted one or more sensory neuron subtypes, as suggested by our earlier transcriptomic analysis. Immunohistochemical markers for sensory neurons, such as calcitonin gene related peptide (CGRP) or isolectin B4 (IB4) for non-peptidergic nociceptors, are liable to change after nerve injury in rodents [73,74]. We therefore used genetic labelling to preserve the identity of sensory neuron subtypes 3 days *in vitro* by expressing the fluorescence reporter TdTomato (TdTom) under the control of one of several canonical marker genes [41]. DRG neurons from *Mrgprd*-expressing mice (a marker for putative mechanosensitive non-peptidergic sensory neurons) [75], showed a high degree of binding by NKG2D-Fc in TdTom+ neurons (**Fig. 4A**), with receptor density highest near the terminals of labelled neurites (**Fig. 4B**), compared to cell bodies (**Fig. 4C**). On the other hand, DRG neurons of the *Th*-expressing (tyrosine hydroxylase expressing C-low threshold mechanoreceptors) [69,76] showed little to no immunoreactivity in TdTom+ (**Fig. 4D**) compared to TdTom-negative neurons (**Fig. 4E**).

**Figure 4.**
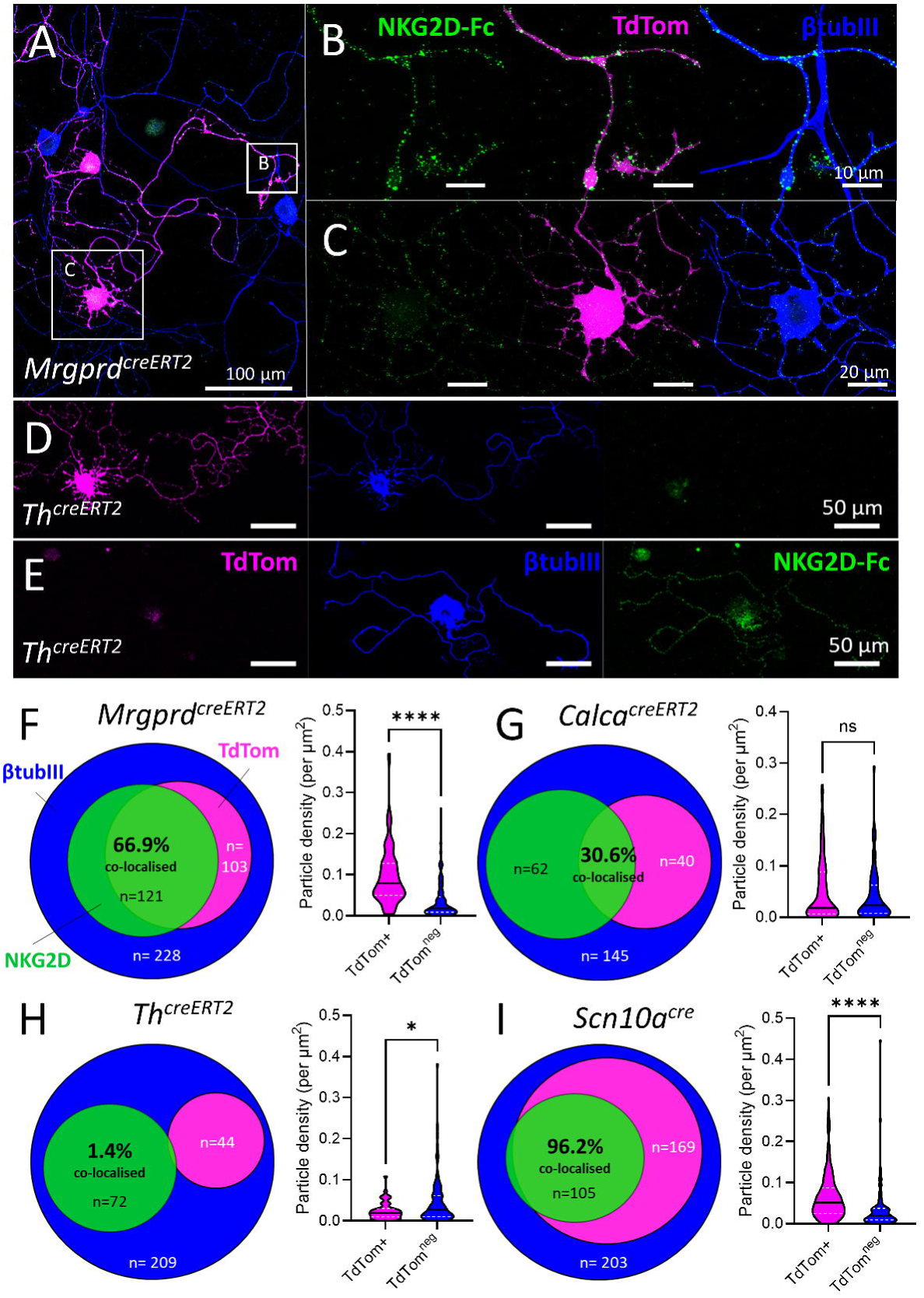
NKG2D receptor binding is enriched among non-peptidergic nociceptive neurons. DRG neuron were cultured for 3 days prior to live cell labelling with NKG2D-Fc. A) Low magnification confocal image of DRG neurons from *Mrgprd-cre^ERT2^;TdTomato* mice 3 days *in vitro*. **B, C)** High magnification images of insets in (A). Note density of NKG2D-Fc receptor binding to neurite terminals (B) compared to cell body (C) of TdTom+ neuron. **D, E**) Low magnification confocal images of DRG neurons from *Th-cre ^ERT2^;TdTomato* mice 3 days *in vitro*. Note lack of NKG2D-Fc receptor binding to neurites of TdTom+ neuron (D) compared to TdTom^neg^ neuron (E). Scale bars as indicated. **F-I**) Quantification of NKG2D receptor binding to the neurites of DRG neurons from four different genetic sensory neuron lineages. Venn diagrams illustrate the proportions of DRG neurons lineages displaying NKG2D binding assessed by manual counting. Violin plots illustrate NKG2D receptor particle density per µm2 of neurite area per image. F) *Mrgprd cre^ERT2^;TdTomato* (non-peptidergic) sensory neurons. Manual counting: n=228 neurons from n=3 mice; 2 male, 1 female. Automated image analysis: TdTom^+^ neurites, n=109 images; TdTom^neg^ neurites, n=129 images; ****p=<0.0001, Kolmogorov-Smirnov test (D=0.5713). **G**) *Calca*-*cre^ERT2^;TdTomato* (peptidergic) sensory neurons. Manual counting: n=145 neurons from n=5 mice; 1 male, 4 female. Automated image analysis: TdTom^+^ neurites, n=67 images; TdTom^neg^ neurites, n=95 images; ns, p=0.5463, Kolmogorov-Smirnov test (D=0.1274). **H**) Tyrosine hydroxylase *(Th)-*lineage (C-LTMR) sensory neurons. Manual counting: n=209 neurons from n=4 mice; 3 male, 1 female. Automated image analysis: TdTom^+^ neurites, n=51 images; TdTom^neg^ neurites, n=100 images; *p=0.0472, Kolmogorov-Smirnov test (D=0.2355). **I**) *Scn10a*-lineage (Nav1.8 nociceptive) sensory neurons. Manual counting: n=203 neurons from n=4 mice; 2 male, 2 female. Automated image analysis: TdTom^+^ neurites, n=162 images; TdTom^neg^ neurites, n=128 images; ****p=<0.0001, Kolmogorov-Smirnov test (D=0.4310). βtubIII counterstain, *blue*. Median and quartiles represented within violin plots as black and white dotted lines, respectively.

Using both blinded manual and automated quantification of NKG2D-Fc labelling of TdTom-positive and negative neurons from each genetic lineage, we observed the greatest enrichment in *Mrgprd*+ neurons (**Fig. 4F**), followed by *Calca*-+ neurons (encoding peptidergic marker CGRP) (**Fig. 4G**), which showed a partial overlap between NKG2D-Fc binding and TdTom+ labelling (**Supp. Figures 3D-K**). Super-resolution imaging of revealed the high density of receptor-bound puncta along the surface of TdTomato-labelled neurites from a *Mrgprd*+ neuron after 3 days *in vitro* (Supplementary Video 1). No significant NKG2D-Fc labelling was observed in *Th*-expressing DRG (**Fig. 4H**). Almost all NKG2D-Fc labelled neurons could be identified by expression of *Scn10a*, encoding the sensory neuron specific voltage-gated sodium channel, Na_v_1.8 (**Fig. 4I**). Similarly, NKG2D-Fc binding was enriched on nociceptive *Trpv1*-lineage neurons (**Supp. Figures 4A, C**) [77], but not *Thy1*-lineage neurons, which label predominantly medium and large diameter DRG neurons [78] (**Supp. Figures 4B, D**).

### Regulation of NKG2D ligands in human sensory neurons

We investigated whether our findings from mice may have any relevance to cytotoxic neuro-immune interactions between human NK cells and sensory neurons by analysing the expression of known NKG2D ligand genes *MICA*, *MICB*, *ULBP1*, *ULBP2*, *ULBP3*, *ULBP4* (*RAET1E*), *ULBP5* (*RAET1G*), and *ULBP6* (*RAET1L*) [31] in published human tissue RNA sequencing datasets.

Findings from two independent bulk RNA sequencing datasets indicated *MICA* (encoding MHC class I polypeptide–related sequence A) as the highest enriched NKG2D ligand gene in primary human DRG tissue and after cell culture [56] (**Supp. Figure 5A**). There was no difference between *MICA* levels in donors with and without pain [57] (**Supp. Figure 5B**). A different pattern was observed in higher resolution single DRG soma (**Fig. 5A**) and spatial (Visium) datasets [58] (**Fig. 5B**) with *ULBP2* in particular featuring predominantly among nociceptive neuron subsets. *MICA*, *MICB*, *ULBP1*, *ULBP3*, *RAET1E*, *REAT1G* and *RAET1L* were inconsistently detected across datasets, being expressed at low level or not detectable (**Fig. 5A, B**). Overall, these findings indicate that NKG2D ligands are expressed in human DRG, likely by nociceptive neurons.

**Figure 5.**
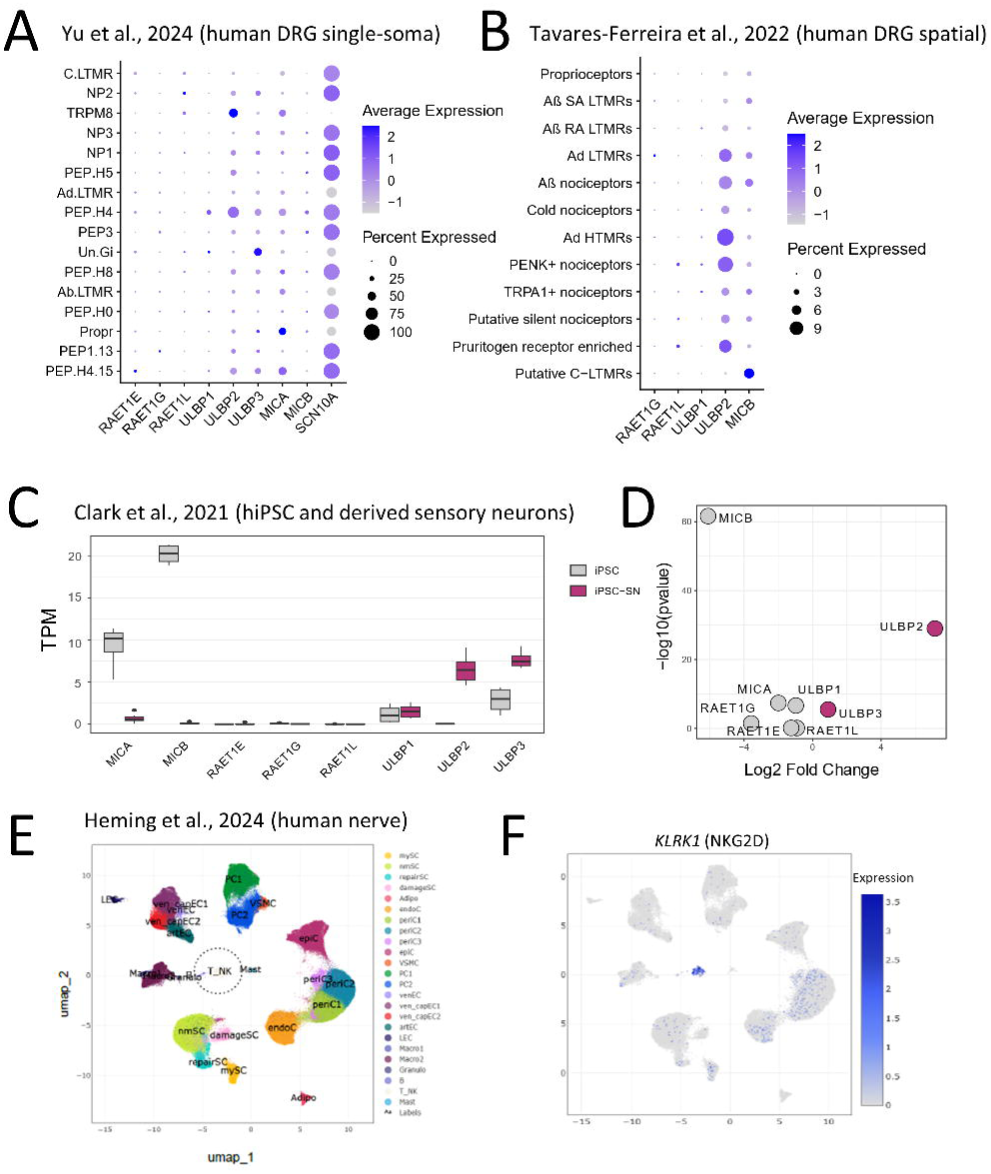
Expression of NKG2D ligands and receptor in human nerve tissues and iPSC models. **A)** Single-soma RNA sequencing of six lower thoracic and lumbar DRG of three human donors (Data from Yu et al., 2024). **B)** Spatial (‘Visium’) RNA sequencing of human lumbar DRG tissues collected from four male and four female organ donors (Data from Tavares-Ferreira et al., 2022). *RAET1E*, *ULBP1* and *MICA* were not detected. **C)** Bulk RNA sequencing of hiPSC and hiPSCd sensory neurons. Log-fold enrichment of NKG2D receptor ligand genes during differentiation of sensory neurons from hiPSC (including lines SFC-AD2-01 (synonym AD2-1, termed ‘AD2’ throughout the study) and SFC-840-03-03 (synonym AH017, termed ‘840’ throughout the study). **D)** *ULBP2* and *ULBP3* show the greatest change in transcript levels (transcripts per million, TPM) after sensory neuron differentiation (Data from Clark et al., 2021). **E)** Identification of 23 cell populations including NK/T cells by analysing the published human sural nerve snRNA-seq data (Data from Heming et al., 2024). **F**) Enrichment of *KLRK1* in NK/T cell subset.

We next compared expression of the same NKG2D ligand genes in human induced pluripotent stem cells (hiPSC) as an approximation of human sensory neurons, in a dataset that included lines ‘840’ and ‘AD2’ [60]. Differential gene expression analysis between hiPSC and differentiated sensory neurons identified *ULBP2* as the most upregulated NKG2D ligand gene, while *MICA* and *MICB* were the genes most downregulated after differentiation to sensory neurons (**Fig. 5C, D**).

Finally, we reanalysed recently published datasets of human neural tissue to understand the potential interaction with NKG2D expressed by localised immune cells. By extracting and re-clustering immune cells present in human DRG tissue [11] (**Supp. Figure 5C**), we identified 13 immune subsets, including one (cluster 9) that expressed several of the 13-gene signature that is characteristic of human NK cells [79] (**Supp. Figure 5D** and was enriched for *KLRK1*, the gene encoding NKG2D (**Supp. Figures 5E, F**). *KLRK1* was also expressed by NK/T cells subsets detected in human sural nerve tissue isolated from patients with polyneuropathy [80] (**Fig. 5E, F**), placing the NKG2D receptor in the same tissue context as its ligands expressed by pathological sensory neurons.

### Functional human NK-sensory neuron interactions

To investigate the functional relevance of NKG2D ligand gene expression we differentiated hiPSC into ‘nociceptor-like’ sensory neurons using a well-characterised protocol [45,46,81,82]. When matured *in vitro* for over 40 days, hiPSCd sensory neurons display characteristic clusters of cell bodies (neuronal soma), with extending BtubIII+ axons/neurites (**Fig. 6A, B**). To investigate if human sensory neurons also regulate NKG2D ligands after injury, and to replicate the injury state of primary cultured mouse DRG, we induced axonal injury in hiPSCd sensory neurons *in vitro* using 355nm laser ablation (**Fig. 6E**). One week after axotomy axons re-grew across the midline of the ablation site (**Fig. 6F** and **Supp. Figure 6A**). Cultures were treated with recombinant human NKG2D-Fc receptors or Fc control proteins (2 µg/ml) 7 days after injury or control (no ablation) and axonal binding was quantified by an automated image acquisition and analysis pipeline (**Supp. Figure 6A**). We observed a non-significant trend of NKG2D binding to uninjured hiPSCd sensory neurons compared to Fc only (**Fig. 6C, D** and **Supp. Figure 6B**). By 7 days after laser axonal ablation human NKG2D receptor binding was significantly higher compared to Fc control (**Fig. 6G, H** and **Supp. Figure 6C**).

**Figure 6.**
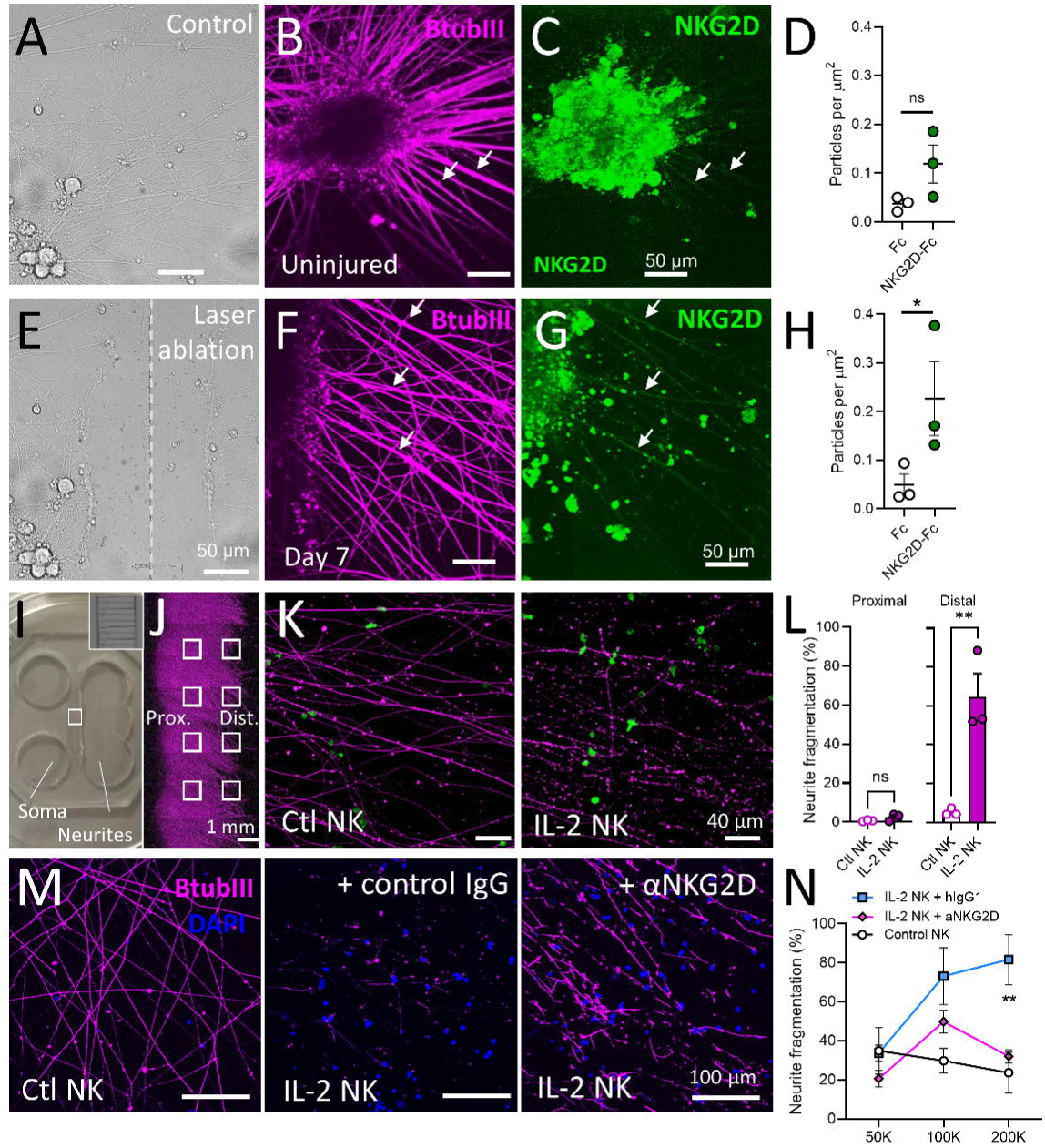
Human iPSC-derived sensory neurons upregulate NKG2D ligands in response to axonal injury and are targets for IL-2 primed human natural killer cells. **A)** Bright-field image of uninjured iPSC-derived sensory neurons. **B)** βtubIII (*magenta*) and **C)** NKG2D-Fc (*green*) immunolabelling of iPSC sensory neurons (‘840’ line) without injury. **D)** Quantification of NKG2D-Fc receptor particle density on βtubIII+ axons compared to Fc-only controls. **E)** Bright-field image of iPSC-derived sensory neurons after axonal injury by laser ablation. **F**) βtubIII (*magenta*) and **G**) NKG2D-Fc (*green*) immunolabelling of iPSC sensory neurons (‘840’ line) 7 days after laser ablation. Arrows highlight axons labelled with NKG2D. **H**) Quantification of NKG2D-Fc receptor particle density on βtubIII+ axons compared to Fc-only controls. n=3 experimental repeats, 2 wells per group, per repeat. Two-way ANOVA, NKG2D versus Fc: F_(1, 8)_ = 8.542, *p=0.0192. Sidak’s multiple comparison test: *p=0.0442. **I**) Photo of microfluidic hiPSCd sensory neuron cultures. Sensory neuron precursors (i.e. neuronal soma) seeded in left hand chamber. Axons grow through microfluidic channels into neurite chamber on right hand side. *Inset*) High magnification of microfluidic channels. **J**) βtubIII immunolabelling of iPSC sensory axons in neurite compartment. Eight regions of interest (*white squares*) were sampled for proximal and distal axon/neurite quantification. **K**) Example ROI of distal axons/neurites from microfluidic devices treated with freshy thawed (control, *left*) or IL-2 primed human NK cells (*right*). NK cells in *gre*en. **L**) Quantification of axon/neurite fragmentation. n=3 experimental repeats (6 microfluidic devices; n=8 ROI per device) with NK cells from three different healthy donors; sensory neurons derived from iPSC donor ‘AD2’. *Proximal* axon fragmentation ns, p=0.1610 (t=1.717); *Distal* axon fragmentation **p=0.0079 (t=4.931), unpaired t test. **M**) Example ROI of distal axons/neurites from microfluidic devices treated with freshy thawed (unstimulated), IL-2 stimulated NK cells incubated with IgG1, IL-2 stimulated NK cells incubated with anti-NKG2D or not treated with NK cells. NK cells in blue (DAPI), axons in magenta (BtubIII). **N**) Microfluidics with different number of NK cells (50×10^3^, 100×10^3^, 200×10^3^) were included for quantification of axon/neurite fragmentation. n=3 experimental repeats (3 microfluidic devices; n=10 ROI per device) with NK cells from three different healthy donors; sensory neurons derived from iPSC donor ‘840’. Multiple comparisons were analysed by two-way ANOVA with Šídák correction. Adjusted p=0.0106 for comparison between microfluidics with unstimulated NK cells and IL-2-stimulated NK cells with IgG1, NK=100×10^3^; adjusted p=0.0009 for comparison between microfluidics with unstimulated NK cells and IL-2-stimulated NK cells with IgG1, NK=200×10^3^; **adjusted p=0.0036 for comparison between microfluidics with IL2-stimulated NK cells incubated with IgG1 and anti-NKG2D, NK=200×10^3^. Scale bars as indicated.

We next asked whether NKG2D receptor ligand expression on human sensory axons had any consequences for human NK cell interactions. To address this possibility, we adapted our previous murine NK-sensory neuron co-culture platform [44] to develop a humanised co-culture system combining human iPSC-derived sensory neurons (line ‘AD2’) grown in microfluidic devices (**Fig. 6I and J**) with human NK cells enriched from the blood of healthy donors by magnetic associated cell sorting (MACS) (**Supp. Figure 7A, B**). Flow cytometry of NK cells showed higher levels of NKG2D (**Supp. Figure 7C-E**), as well and perforin and granzyme B (**Supp. Figure 7F-I**), after priming with human interleukin-2 (IL-2) (1000 U/ml for 2 days) compared to untreated controls, confirming a cytotoxic gain-of-function. Addition of NK cells to the neurite compartment of microfluidic devices hiPSCd-sensory neurons resulted in fragmentation of distal axons by IL-2 primed NK cells (**Fig. 6K, L**), reminiscent of the effects in murine NK-DRG co-cultures [18,44].

Our findings of possible low-level binding of NKG2D receptors to ‘uninjured’ hiPSCd sensory neurons prompted us to investigated whether human NKG2D receptor-ligand interactions may play a role in this neurodegenerative interaction, this time using a different hiPSC cell line (‘840’) (**Fig. 6M**). Pre-treatment of the IL-2-primed human NK cells with an NKG2D blocking antibody led to a reduction in axon fragmentation compared to isotype control (**Fig. 6N**), suggesting that NKG2D receptor ligands may contribute to the degeneration of human sensory axons by primed NK cells.

## Discussion

In this study, we used a combination of transcriptomic and targeted protein analysis to identify surface expression of NKG2D ligands by injured nociceptive neurons in mice and humans. These findings have three important implications: 1) They provide a molecular basis for understanding the neuropathic pain-resolving capacity of NK cells by targeting a subset of nociceptive nerve fibres known to be crucial for the neuropathic phenotype in mice after traumatic nerve injury. 2) This reinforces the capacity for pathological nociceptors to modulate the local immune microenvironment through the expression of ligands for potent immune receptors. 3) The specific ligands identified in both mice and humans in this study could serve not only as unique markers of pathological nerve injury, but also targets for novel diagnostic methods and ultimately therapeutic intervention.

### Molecular identity of murine DRG neurons expressing NKG2D ligands

In healthy tissues, NKG2D ligands are expressed at low or non-detectable levels. We therefore adopted several complementary approaches to triangulate the genetic and functional identity of mouse sensory neurons expressing NKG2D ligands after nerve injury.

We show that *Raet1e* is most likely the predominant isoform expressed by injured nociceptive DRG neurons in C57BL/6 mice (**Fig. 1**). *Raet1e* shows 25-fold higher affinity for the NKG2D receptor than *Raet1d* [83], suggesting that the former is likely to be the more functionally relevant neuro-immune interaction in the context of peripheral nerve injury [29]. We cannot exclude the possible contribution from myeloid cells (e.g. macrophages) to the detection of *Raet1d* and *Raet1e* transcripts in whole DRG tissues [84]. Transcripts for another NKG2D ligand, *Mult1* (*Ulbp1*) was also upregulated in DRG by nerve injury, but was detected in only two out of 38 successfully collected single DRG neurons (sham or L5x).

Although a number of NKG2D ligand genes appear in publicly available RNA sequencing datasets of mouse DRG, *Raet1e* was not reliably identified as differentially expressed by nerve injury [42,85], or in any specific subset(s) of DRG neurons [67,69,86]. To overcome the potential limitations in sequencing depth in single cell datasets, we analysed transcripts from pooled DRG neurons enriched for neuronal subsets [41]. This approach not only confirmed regulation of individual NKG2D ligand genes *Raet1e* and *Mult1* (*Ulbp1*) by nerve injury, but also provided clues to the subset involved (*Mrgprd*-expressing, non-peptidergic neurons) (**Fig. 1**).

Using highly sensitive nested PCR, we were able to confirm *Raet1* genes in individual DRG neurons. The ubiquitous expression of *Runx1* and high prevalence of *Scn10a* among single cells collected for PCR again reinforced their identity as either nociceptors (**Fig. 2**) [87] or C-LTMRs [70], while only around 50% were positive for Trka, which is transcriptomic marker for peptidergic neurons [71]. A very recently published high-quality, integrated scRNA-seq atlas of cells in mouse DRG revealed *Raet1e* and *Mult1* (*Ulbp1*) to be expressed by non-peptidergic nociceptors and C-LTMRs, respectively [70]. While this finding is in keeping with the co-detection of *Mult1* with transcripts for *Runx1* and *Scn10a* by single-cell PCR (both also expressed by C-LTMRs) [88], the detection of a transcript for an NKG2D ligand gene in C-LTMRs stands in contrast to our live cell assays where no receptor binding was observed (see below).

Interestingly, there was no difference in the proportion of DRG neurons expressing *Raet1* after L5x injury or sham surgery, suggesting that the increase in transcript we observed by cell culture *in vitro,* or after injury *in vivo,* occurred within the same population of neurons. We also did not observe any noticeable sex-differences in our data suggesting that NKG2D ligand regulation is a likely feature of nerve injury in female as well as male mice.

### NKG2D as a functional biomarker nociceptive neuro-immune interactions after injury

To overcome the potential discrepancy between transcript and protein levels, and to better understand the dynamics of cell surface expression, we adopted a live cell-based assay approach to identify NKG2D ligands [27]. Live cell assays have proved to be exquisitely sensitive to identify surface binding to neuronal antigens in heterologous systems [89], as well as live neurons [90]. We revealed NKG2D receptor binding to discrete puncta along the neurites of DRG (**Fig. 3**) that mirrored the timeline of transcriptional upregulation in culture (**Fig. 1**). The apparent enrichment of NKG2D receptor binding to distal neurites is reminiscent of Raet1 protein labelling in chronically injured sciatic nerve *in vivo* [18] and provides an anatomical framework for potential cytotoxic neuro-immune interactions at the distal nerve injury site [91]. Whether the peripheral compartmentalisation of receptor ligands in DRG neurons occurs due to axonal transport, or local RNA translation, remains to be investigated.

The process of culturing DRG neurons – effectively an axotomy – has long been known to affect their biophysical properties [74]. The parallel upregulation of *Raet1e* and *Atf3* emphasise that DRG neurons enter an injury-like state *in vitro* (**Fig. 1B**). The use of fluorescent reporter mouse lines allowed us to overcome the potential loss of biochemical markers induced by the cell culture process. Consistent with our transcriptional analysis of NKG2D ligands, soluble NKG2D receptors bound exclusively to neurons of a nociceptive lineage identified by developmental expression of *Scn10a* (**Fig. 4**) and *Trpv1*. Conversely, the predominant lack of recognition of *Thy1*-lineage fibres suggests that large diameter fibres are not a target for NKG2D recognition [78]. Among nociceptors, we observed preferential binding to neurites of the non-peptidergic (*Mrgprd*-expressing) subpopulation of DRG neurons, with a smaller proportion recognising peptidergic (*Calca*-lineage) neurons (**Fig. 4**). This finding is consistent with the single cell PCR where around 50% of neurons expressing *Raet1* were also positive for peptidergic marker TrkA (**Supp. Figure 2**) and with data from bulk sequencing of DRG neurons from the same mouse lines after nerve injury (**Fig. 1**) [41].

Interestingly, *Th*-lineage low-threshold mechano-sensitive C-fibres were not among those to bind soluble NKG2D receptors (**Fig. 4**). This finding that stands in contrast to the latest integrated scRNA-seq atlas of mouse DRG neurons, which revealed the majority of *Ulbp1* (*Mult1*) expressing DRG to be C-LTMRs [70]. There are several possible explanations for this discrepancy, which require further investigation: 1) *Ulbp1* (*Mult1*) transcripts may not be functionally expressed as cell surface ligands. 2) Ulbp1 ligands may not be recognised by soluble NKG2D receptors. 3) Ulbp1 ligands may not be accessible to receptor binding at the cell surface of C-LTMRs.

High and low-threshold C-fibres are both functionally and transcriptionally distinct [69,92], therefore neuro-immune interactions between either of these two neuronal subsets is likely to have differing pathophysiological relevance for somatosensation. Unmyelinated nociceptors, and the non-peptidergic subset in particular, have been implicated in neuropathic pain in mice after nerve injury or peripheral neuropathy [75,93–95]. A population of small diameter nociceptive neurons expressing *Mrgprd* (i.e. non-peptidergic) are thought to undergo cell death after traumatic nerve injury [96]. Whether the appearance of functional NKG2D ligands at the soma of these DRG neurons after injury renders them susceptible to immune-mediated killing remains an intriguing possibility.

Aberrant regeneration of unmyelinated sensory afferents after insult or injury has well-documented pathological consequences for somatosensation. Collateral sprouting of unmyelinated high-threshold nociceptors into Meissner corpuscles of denervated territories after nerve injury drives mechanical hypersensitivity [97]. Similarly, re-wiring of Merkel cells by non-peptidergic nociceptors thought to underlying chronic itch in a model of dry skin [98]. Conversely, sprouting is not observed by low threshold Aβ fibres or C fibres [99], which are thought to undergo gain-of-function within adjacent uninjured territories after nerve-injury.

Preferential expression of NKG2D ligands by putative pathological Mrgprd+ high-threshold mechanoreceptors [75,95] as they reinnervate peripheral tissues may help explain the efficacy of NK cell stimulation to resolve neuropathic mechanical hypersensitivity after partial crush nerve injury [18], a preclinical model of neuropathic pain in which axons are not restricted from regenerating [100]. Whether other sensory modalities, such as pathological itch, are affected by NK cell-sensory neurons interactions remains to be explored.

### Regulation of NKG2D ligands by pathological neurons

NKG2D ligands are typically absent from otherwise healthy cells, but become upregulated in in response to tumorigenic transformation, infection, as well as cellular stressors. Expression of *Raet1e* has been link to activity of E2F transcription factors that control cell cycle and are commonly dysregulated in cancer [101]. Infection with murine cytomegalovirus (MCMV) induces *Raet1e* expression by switching off the transcription suppressor histone deacetylase inhibitor 3 (HDAC3) [102]. DNA damage response is another important factor driving NKG2D ligand upregulation, especially mouse *Raet1* and *Mult1* [103]. Importantly, *Raet1* and *Mult1* can be regulated by distinct pathways [104], which may explain the discrepancy in expression we observed between these two ligands in DRG neuron subsets.

Within the healthy adult nervous system, *Raet1* gene expression is found during embryogenesis in the brain [105] and DRG [18], and as well as rare niches of neurogenesis in the adult brain [106] suggesting a link to neuronal development. In addition to nerve injury [18], other examples of NKG2D ligand regulation in mouse models of neuropathology include cerebral ischaemia [107], experimental autoimmune encephalitis [84] and amyotrophic lateral sclerosis [108]. Although generally regarded as post-mitotic, adult DRG neurons respond to nerve injury by activation of a regeneration associated gene programme that results in a genetic de-differentiation [12]. Recent evidence suggests DRG in mice display a senescence-like state reminiscent of cell cycle arrest after both injury and aging that appears to be associated with a hyperexcitable neuronal phenotype [109,110]. This raises the question of whether ageing, as well as other potential initiators of peripheral neuropathy - such as diabetes [111,112] and chemotherapy [113,114] - may involve the regulation of NKG2D ligand expression by nociceptive neurons.

The de novo expression of surface ligands for NKG2D after nerve injury [18] reinforces the idea that pathological nociceptor subtypes, as identified in this paper, are potential targets for immune surveillance [115] and therapeutic intervention [19,116]. How NKG2D ligand expression by pathological nociceptive sensory neurons may regulate the local immune microenvironment of innervated organs and barrier tissues, and how this may reciprocally affect nociceptor function as with other peripheral neuro-immune interactions [117], remains to be explored.

### Translation of NK-sensory neuron interactions to humans

We identified transcripts for a number of known NKG2D ligand transcripts in published human DRG and hiPSCd sensory neuron datasets. Among the most consistently identified transcript at near-single cell resolution was *ULBP2* (**Fig. 5**). We additionally identified that human NKG2D receptor-binding to nociceptor-like human iPSC-derived sensory neurons significantly increases after axon ablation (**Fig. 6**), indicating the upregulation of cell surface NKG2D ligands after nerve injury. These results echo earlier finding of ULBP expression by intraepidermal nerve fibres in people with fibromyalgia [118], strengthening the potential functional role that NKG2D-ligand interactions may play between NK cells and human nociceptive neurons in peripheral nerve pathology. The identification of *KLRK1*-expression on non-neuronal cells with a gene profile consistent with cytotoxic NK and T lymphocytes within human peripheral nervous tissues (**Fig. 5**) further reinforces the feasibility of NKG2D receptor-ligand interactions between immune cell populations and peripheral sensory neurons in a clinical context.

Compared to mice, the distinction between subtypes of human nociceptors is less clear. Human DRG show a greater overlap in expression of *CALCA*+ and *P2X3*+, which in mice are markers traditionally associated with peptidergic and non-peptidergic neurons, respectively [119]. There is also higher prevalence of human DRG neurons expressing classical peptidergic markers CGRP and TRPV1 [120]. While the human iPSC differentiation protocol we employed is well-validated [46] it must be remembered these cells offer only an approximation of human nociceptive DRG neurons [121]. Alternative differentiation protocols, which result in a greater number of TRPV1 and CGRP-positive neurons [82] may provide more translatable findings.

The difference in findings from bulk RNA sequencing datasets from whole tissue and cultured DRG, which had a higher prevalence of *MICA* and *MICB* transcripts, could indicate their expression by non-neuronal cells, including lymphocytes and satellite glia, within human DRG [11,122]. The interaction between NKG2D and its ligands expressed on non-neuronal cells in the context of peripheral nerve injury remains to be explored.

Consistent with previous findings in mice [18], degeneration of hiPSCd sensory axons by primary human NK cells was prevented by an NKG2D blocking antibody (**Fig. 6**). This effect was observed in the absence of axonal injury, which suggests hiPSCd sensory neurons may either retain developmental expression of NKG2D ligands from their hiPSC identity, or that the differentiated neurons exist in a basal ‘stress-like’ state. Replication of the experiment with live primary cultured human DRG would help provide a definitive answer. While comparing our observations of sensory neurons in the two species should be approached with caution owing to their developmental origins, the enhanced binding of NKG2D we observed to hiPSCd sensory axons after laser axon ablation suggests that axotomy alone is sufficient to induce an injury-like state. Further work is required to validate the characteristics of hiPSCd sensory neurons as a model of injury-induced neuropathology.

Diagnosis of neuropathic pain specifically within chronic pain conditions is often challenging. Questionnaire-based screening tools such as DN4 (douleur neuropathique 4 questions), NPSI (Neuropathic Pain Symptom Inventory) and PainDETECT [123] are helpful, but still rely on a detailed and labour-intensive clinical assessment to identify a lesion or disease of the somatosensory nervous system [124]. There has therefore been a recent focus on obtaining objective biomarkers to better understand the subjective experience of pain [125]. Composite factors can be applied at the population level [126] but diagnosis at the patient level remains challenging. The search for biochemical biomarkers of nerve injury has therefore typically focussed on easily accessible analytes in the peripheral blood. For example, a recent systematic review highlighted significantly higher levels of neurofilament light chain in cases of peripheral neuropathy compared to controls [127]. A cell surface-expressed protein biomarker that is found at low levels in ‘healthy’ individuals and upregulated by peripheral nerve injury - as identified here in the form of Raet1e (mouse) and ULBP2 (human) - could pave the way for identifying and intervention in nerve damage via non-invasive imaging or radiolabelling techniques.

### Limitations

This study was performed in mice of both sexes on a C57BL/6J background using animals sourced from suppliers at three geographical locations (UK, South Korea and US). While the sex and supplier variables enhance the external validity of our findings, the use of a single genetic background, which are known to express different NKG2D ligands compared to other strains such as Balb/c [33], is a significant limitation. An examination of NKG2D ligands after nerve injury in other strains of mice would strengthen the claims of the evolutionary conservation of this particular neuro-immune interaction.

Several of the cre recombinase-driven reporter mouse lines used, including *Trpv1*-cre, restricts our interpretation to sensory neurons of a particular genetic lineage [38]. This was overcome by the use of tamoxifen inducible cre lines, in which Cre recombinase is fused to a mutant oestrogen ligand-binding domain (ERT2), allows for developmental control of reporter transgene (TdTomato) expression [41]. However, we cannot exclude a potential discrepancy between lineage markers and functional identity of sensory neurons that express NKG2D ligands in the adult mouse, nor a potential effect of tamoxifen itself on driving cellular stress in addition to the effect of injury [128]. These findings therefore emphasise the importance of cross-validating potential receptor-ligand interactions at multiple molecular and cellular levels, especially when genes expressed even at low levels can have powerful neuro-immune modulatory functions.

The small sample size for single cell PCR experiment (18-20 *Gapdh+Advillin+* DRG neurons per group) limits interpretation of the minimal detection of Mult1. The high proportion of small diameter neurons among those collected also emphasises the sample bias inherent in DRG neuron culture [44].

We focussed our receptor binding analysis on the neurites of DRG neurons *in vitro*. The purpose of the soma extraction step in our analysis was to avoid the confound of dead/dying cells, which non-specifically attract recombinant protein and antibody binding. Although we provide anecdotal evidence for NKG2D ligand expression predominantly at distal neurites, there remains a potential for functional receptor expression at DRG soma. Further work investigating ligand dynamics will help shed light on their subcellar distribution, and whether cytotoxic neuroimmune interactions at the soma may have consequences for neuron viability after nerve injury [96].

Our use of primary human NK cells inevitably results in a mismatch between donor and recipient human leukocyte antigen (HLA), which may affect the influence the neuro-immune interaction we observe *in vitro* in a way that would not occur within an individual with nerve injury. Exposure of host immune system to tissue from an HLA-mismatched individual – as occurs in organ transplants – can result in graft-versus-host disease (GvHD) that is typically the result of alloreactive T cell activity. Although the role of NK cells is more complex, which in some cases are thought to offer protection from GvHD [129], we cannot exclude the possibility of alloreactive NK cells responding to HLA-mismatch with the iPSC lines [130] contributing to the axon degeneration we observed after IL-2 priming.

## Conclusion

The functional expression of NKG2D receptor ligands at the cell surface of unmyelinated nerve fibres may represent an anatomically restricted marker of injury, or other nerve pathology. De novo display of ligands for receptors of immune surveillance on non-peptidergic nociceptors in particular, is likely to result in reciprocal neuro-immune interactions that influence somatosensation and pain, as well as the local immune microenvironment, in injury or disease.

## Data availability

All published datasets used in this study are cited in the results and figure legends. Flow cytometry data are available from Zenodo [131]. Other summary data are available upon reasonable request to the Corresponding author.

## Code availability

Detailed and annotated scripts for analysing NKG2D binding to neurites of murine DRG neurons and human iPSC-derived sensory neurons, as well as script for calculating the fragmentation of axons, are available via GitHub (see Methods).

## Author contributions

Conceptualization, AJD

Methodology, AJD, SW, AMB, NY, LSB, YKL, SWS

Software, HCK Validation, SW

Formal analysis, AJD, SW, AMB, GB, YKL

Investigation, AJD, SW, AMB, YKK, NY, XW, LSB, RGC, YKL, SWS

Resources, SR, DLB, MC

Data Curation, SW, AMB, HCK, GB

Visualization, AJD, SW, YKL

Writing - Original Draft, AJD, SW, SWS

Writing - Review & Editing, AJD, SW, SWS, YKL, AMB, SBO, DLB

Supervision, AJD, SBO, MC, DLB

Funding acquisition, AJD, AMB, SBO, DLB, SR

Project administration, AJD

## Acknowledgements

The authors would like to thank Dr Chul-Kyu Park for assistance with nested primer design, Dr Demetrio Labate and Dr Cihan Bilge Kayasandik for guidance and implementation of the Directional Ratio method, and Dr Alan Wainman for assistance with the SoRa confocal microscopy. We thank Professors Pao-Tien Chuang, John Wood, and Pilhan Kim for generously providing some of the transgenic mice used in this study. We thank Dr Jussi Kupari and Professor Patrik Ernfors for discussion and generously sharing data prior to publication. We also thank James Barker and the rest of the RIVER Working Group for feedback during the drafting of this manuscript.

## Abbreviations

ATF3: Activating Transcription Factor 3
B2m: Beta-2 microglobulin
C-LTMRs: C-low threshold mechanoreceptors
CGRP: calcitonin gene related peptide
DRG: Dorsal root ganglion
H60: Histocompatibility 60
hiPSCd: Human induced pluripotent stem cell-derived
IB4: Isolectin B4
IL-2: Interleukin-2
L5x: 5^th^ lumbar spinal nerve transection
Mult1: Murine UL16-binding protein-like transcript 1
NGF: Nerve growth factor
NK: Natural killer
NKG2D: Natural killer group 2D
Raet1: Retinoic acid early transcript 1
RTx: Resiniferatoxin
SNI: Spared nerve injury
TRKA (NRTK1): Neurotrophic Receptor Tyrosine Kinase 1
TRPV1: Transient Receptor Potential Vanilloid 1
Ulbp: UL16-binding protein-like

## References

[1]. Basbaum AI, Bautista DM, Scherrer G, Julius D. Cellular and molecular mechanisms of pain. Cell. 2009;139(2):267–84.

[2]. Stoll G, Muller HW. Nerve injury, axonal degeneration and neural regeneration: basic insights. Brain Pathol. 1999;9(2):313–25.

[3]. Costigan M, Scholz J, Woolf CJ. Neuropathic pain: a maladaptive response of the nervous system to damage. Annu Rev Neurosci. 2009;32:1–32.

[4]. Knezevic NN, Cicmil N, Knezevic I, Candido KD. Discontinued neuropathic pain therapy between 2009-2015. Expert Opin Investig Drugs. 2015;24(12):1631–46.

[5]. Kingwell K. Nav1.7 withholds its pain potential. Nat Rev Drug Discov. 2019.

[6]. Soliman N, Moisset X, Ferraro MC, de Andrade DC, Baron R, Belton J, et al. Pharmacotherapy and non-invasive neuromodulation for neuropathic pain: a systematic review and meta-analysis. The Lancet Neurology. 2025;24(5):413–28.

[7]. Finnerup NB, Attal N, Haroutounian S, McNicol E, Baron R, Dworkin RH, et al. Pharmacotherapy for neuropathic pain in adults: a systematic review and meta-analysis. Lancet Neurol. 2015;14(2):162–73.

[8]. Woolf CJ. Capturing Novel Non-opioid Pain Targets. Biol Psychiatry. 2020;87(1):74–81.

[9]. Qi L, Iskols M, Shi D, Reddy P, Walker C, Lezgiyeva K, et al. A mouse DRG genetic toolkit reveals morphological and physiological diversity of somatosensory neuron subtypes. Cell. 2024;187(6):1508–26 e16.

[10]. Rostock C, Schrenk-Siemens K, Pohle J, Siemens J. Human vs. Mouse Nociceptors - Similarities and Differences. Neuroscience. 2018;387:13–27.

[11]. Bhuiyan SA, Xu M, Yang L, Semizoglou E, Bhatia P, Pantaleo KI, et al. Harmonized cross-species cell atlases of trigeminal and dorsal root ganglia. Sci Adv. 2024;10(25):eadj9173.

[12]. Renthal W, Tochitsky I, Yang L, Cheng YC, Li E, Kawaguchi R, et al. Transcriptional Reprogramming of Distinct Peripheral Sensory Neuron Subtypes after Axonal Injury. Neuron. 2020;108(1):128–44 e9.

[13]. Dib-Hajj S, Black JA, Felts P, Waxman SG. Down-regulation of transcripts for Na channel alpha-SNS in spinal sensory neurons following axotomy. Proceedings of the National Academy of Sciences of the United States of America. 1996;93(25):14950–4.

[14]. Jones J, Correll DJ, Lechner SM, Jazic I, Miao X, Shaw D, et al. Selective Inhibition of Na(V)1.8 with VX-548 for Acute Pain. N Engl J Med. 2023;389(5):393–405.

[15]. Jain A, Hakim S, Woolf CJ. Immune drivers of physiological and pathological pain. J Exp Med. 2024;221(5).

[16]. Pinho-Ribeiro FA, Verri WA, Jr., Chiu IM. Nociceptor Sensory Neuron-Immune Interactions in Pain and Inflammation. Trends Immunol. 2017;38(1):5–19.

[17]. Fiore NT, Debs SR, Hayes JP, Duffy SS, Moalem-Taylor G. Pain-resolving immune mechanisms in neuropathic pain. Nat Rev Neurol. 2023;19(4):199–220.

[18]. Davies AJ, Kim HW, Gonzalez-Cano R, Choi J, Back SK, Roh SE, et al. Natural Killer Cells Degenerate Intact Sensory Afferents following Nerve Injury. Cell. 2019;176(4):716–28 e18.

[19]. Kim HW, Wang S, Davies AJ, Oh SB. The therapeutic potential of natural killer cells in neuropathic pain. Trends Neurosci. 2023;46(8):617–27.

[20]. Vivier E, Artis D, Colonna M, Diefenbach A, Di Santo JP, Eberl G, et al. Innate Lymphoid Cells: 10 Years On. Cell. 2018;174(5):1054–66.

[21]. Chan CJ, Smyth MJ, Martinet L. Molecular mechanisms of natural killer cell activation in response to cellular stress. Cell Death Differ. 2014;21(1):5–14.

[22]. Lanier LL. Five decades of natural killer cell discovery. J Exp Med. 2024;221(8).

[23]. Houchins JP, Yabe T, McSherry C, Bach FH. DNA sequence analysis of NKG2, a family of related cDNA clones encoding type II integral membrane proteins on human natural killer cells. J Exp Med. 1991;173(4):1017–20.

[24]. Wensveen FM, Jelencic V, Polic B. NKG2D: A Master Regulator of Immune Cell Responsiveness. Front Immunol. 2018;9:441.

[25]. Lopez-Larrea C, Suarez-Alvarez B, Lopez-Soto A, Lopez-Vazquez A, Gonzalez S. The NKG2D receptor: sensing stressed cells. Trends Mol Med. 2008;14(4):179–89.

[26]. Nomura M, Takihara Y, Shimada K. Isolation and characterization of retinoic acid-inducible cDNA clones in F9 cells: one of the early inducible clones encodes a novel protein sharing several highly homologous regions with a Drosophila polyhomeotic protein. Differentiation. 1994;57(1):39–50.

[27]. Cerwenka A, Bakker AB, McClanahan T, Wagner J, Wu J, Phillips JH, et al. Retinoic acid early inducible genes define a ligand family for the activating NKG2D receptor in mice. Immunity. 2000;12(6):721–7.

[28]. Diefenbach A, Jensen ER, Jamieson AM, Raulet DH. Rae1 and H60 ligands of the NKG2D receptor stimulate tumour immunity. Nature. 2001;413(6852):165–71.

[29]. Carayannopoulos LN, Naidenko OV, Fremont DH, Yokoyama WM. Cutting edge: murine UL16-binding protein-like transcript 1: a newly described transcript encoding a high-affinity ligand for murine NKG2D. J Immunol. 2002;169(8):4079–83.

[30]. Sutherland CL, Chalupny NJ, Cosman D. The UL16-binding proteins, a novel family of MHC class I-related ligands for NKG2D, activate natural killer cell functions. Immunol Rev. 2001;181:185–92.

[31]. Kubin M, Cassiano L, Chalupny J, Chin W, Cosman D, Fanslow W, et al. ULBP1, 2, 3: novel MHC class I-related molecules that bind to human cytomegalovirus glycoprotein UL16, activate NK cells. Eur J Immunol. 2001;31(5):1428–37.

[32]. Lanier LL. NKG2D Receptor and Its Ligands in Host Defense. Cancer Immunol Res. 2015;3(6):575–82.

[33]. Ogasawara K, Lanier LL. NKG2D in NK and T cell-mediated immunity. J Clin Immunol. 2005;25(6):534–40.

[34]. . The-RIVER-Working-Group. Reporting In Vitro Experiments Responsibly – the RIVER Recommendations. MetaArXiv. 2023:1–43.

[35]. Percie du Sert N, Hurst V, Ahluwalia A, Alam S, Avey MT, Baker M, et al. The ARRIVE guidelines 2.0: Updated guidelines for reporting animal research. PLoS Biol. 2020;18(7):e3000410.

[36]. Abraira VE, Kuehn ED, Chirila AM, Springel MW, Toliver AA, Zimmerman AL, et al. The Cellular and Synaptic Architecture of the Mechanosensory Dorsal Horn. Cell. 2017;168(1-2):295–310 e19.

[37]. Olson W, Abdus-Saboor I, Cui L, Burdge J, Raabe T, Ma M, et al. Sparse genetic tracing reveals regionally specific functional organization of mammalian nociceptors. Elife. 2017;6.

[38]. Cavanaugh DJ, Chesler AT, Jackson AC, Sigal YM, Yamanaka H, Grant R, et al. Trpv1 reporter mice reveal highly restricted brain distribution and functional expression in arteriolar smooth muscle cells. The Journal of neuroscience : the official journal of the Society for Neuroscience. 2011;31(13):5067–77.

[39]. Wu S, Wu Y, Capecchi MR. Motoneurons and oligodendrocytes are sequentially generated from neural stem cells but do not appear to share common lineage-restricted progenitors in vivo. Development. 2006;133(4):581–90.

[40]. Madisen L, Zwingman TA, Sunkin SM, Oh SW, Zariwala HA, Gu H, et al. A robust and high-throughput Cre reporting and characterization system for the whole mouse brain. Nat Neurosci. 2010;13(1):133–40.

[41]. Barry AM, Zhao N, Yang X, Bennett DL, Baskozos G. Deep RNA-seq of male and female murine sensory neuron subtypes after nerve injury. Pain. 2023;164(10):2196–215.

[42]. Cobos EJ, Nickerson CA, Gao F, Chandran V, Bravo-Caparros I, Gonzalez-Cano R, et al. Mechanistic Differences in Neuropathic Pain Modalities Revealed by Correlating Behavior with Global Expression Profiling. Cell Rep. 2018;22(5):1301–12.

[43]. Kim YH, Back SK, Davies AJ, Jeong H, Jo HJ, Chung G, et al. TRPV1 in GABAergic interneurons mediates neuropathic mechanical allodynia and disinhibition of the nociceptive circuitry in the spinal cord. Neuron. 2012;74(4):640–7.

[44]. Kim HW, Davies AJ, Oh SB. In Vitro Visualization of Cell-to-Cell Interactions Between Natural Killer Cells and Sensory Neurons. Methods Mol Biol. 2022;2463:251–68.

[45]. Clark AJ, Kaller MS, Galino J, Willison HJ, Rinaldi S, Bennett DLH. Co-cultures with stem cell-derived human sensory neurons reveal regulators of peripheral myelination. Brain. 2017;140(4):898–913.

[46]. Chambers SM, Qi Y, Mica Y, Lee G, Zhang XJ, Niu L, et al. Combined small-molecule inhibition accelerates developmental timing and converts human pluripotent stem cells into nociceptors. Nat Biotechnol. 2012;30(7):715–20.

[47]. Clark AJ. Establishing Myelinating Cocultures Using Human iPSC-Derived Sensory Neurons to Investigate Axonal Degeneration and Demyelination. Methods Mol Biol. 2020;2143:111–29.

[48]. Schindelin J, Arganda-Carreras I, Frise E, Kaynig V, Longair M, Pietzsch T, et al. Fiji: an open-source platform for biological-image analysis. Nat Methods. 2012;9(7):676–82.

[49]. Kayasandik CB, Labate D. Improved detection of soma location and morphology in fluorescence microscopy images of neurons. J Neurosci Methods. 2016;274:61–70.

[50]. DaviesLab-Oxford. (2025) NKG2D-binding-murine-axons-lineage-analysis-pipeline GitHub (Zenodo). 10.5281/zenodo.15921009 (Accessed July 2025)

[51]. DaviesLab-Oxford. (2025) NKG2D-binding-hiPSCdSN-analysis-pipeline GitHub (Zenodo). 10.5281/zenodo.15920869 (Accessed July 2025)

[52]. DaviesLab-Oxford. (2025) Fragmentation-axons-automated-analysis-Fiji GitHub (Zenodo). 10.5281/zenodo.15920937 (Accessed July 2025)

[53]. Schmittgen TD, Livak KJ. Analyzing real-time PCR data by the comparative C(T) method. Nature protocols. 2008;3(6):1101–8.

[54]. Ye J, Coulouris G, Zaretskaya I, Cutcutache I, Rozen S, Madden TL. Primer-BLAST: a tool to design target-specific primers for polymerase chain reaction. BMC Bioinformatics. 2012;13:134.

[55]. Kibbe WA. OligoCalc: an online oligonucleotide properties calculator. Nucleic Acids Res. 2007;35(Web Server issue):W43–6.

[56]. Wangzhou A, McIlvried LA, Paige C, Barragan-Iglesias P, Shiers S, Ahmad A, et al. Pharmacological target-focused transcriptomic analysis of native vs cultured human and mouse dorsal root ganglia. Pain. 2020;161(7):1497–517.

[57]. Ray PR, Shiers S, Caruso JP, Tavares-Ferreira D, Sankaranarayanan I, Uhelski ML, et al. RNA profiling of human dorsal root ganglia reveals sex differences in mechanisms promoting neuropathic pain. Brain. 2023;146(2):749–66.

[58]. Tavares-Ferreira D, Shiers S, Ray PR, Wangzhou A, Jeevakumar V, Sankaranarayanan I, et al. Spatial transcriptomics of dorsal root ganglia identifies molecular signatures of human nociceptors. Sci Transl Med. 2022;14(632):eabj8186.

[59]. Yu H, Nagi SS, Usoskin D, Hu Y, Kupari J, Bouchatta O, et al. Leveraging deep single-soma RNA sequencing to explore the neural basis of human somatosensation. Nat Neurosci. 2024.

[60]. Clark AJ, Kugathasan U, Baskozos G, Priestman DA, Fugger N, Lone MA, et al. An iPSC model of hereditary sensory neuropathy-1 reveals L-serine-responsive deficits in neuronal ganglioside composition and axoglial interactions. Cell Rep Med. 2021;2(7):100345.

[61]. Bhuiyan SA, Renthal W. (2024) Harmonized DRG and TG reference atlas. Painseq (Shinyapp). Available from: https://painseq.shinyapps.io/harmonized_painseq_v1/ (Accessed June 2025)

[62]. Heming M, Hörste GMz. (2024) Single nuclei sural nerve atlas. (Cerebro). Available from: https://osmzhlab.uni-muenster.de/shiny/cerebro_pns_atlas/ (Accessed June 2025)

[63]. Krauter D, Ernfors P. (2025) Mouse DRG Integrated Atlas. Ernforslab (Shinyapp). Available from: https://ernforslab.shinyapps.io/integratedDRGatlas/ (Accessed June 2025)

[64]. Girardi M, Oppenheim DE, Steele CR, Lewis JM, Glusac E, Filler R, et al. Regulation of cutaneous malignancy by gammadelta T cells. Science. 2001;294(5542):605–9.

[65]. Ogasawara K, Benjamin J, Takaki R, Phillips JH, Lanier LL. Function of NKG2D in natural killer cell-mediated rejection of mouse bone marrow grafts. Nat Immunol. 2005;6(9):938–45.

[66]. Takada A, Yoshida S, Kajikawa M, Miyatake Y, Tomaru U, Sakai M, et al. Two novel NKG2D ligands of the mouse H60 family with differential expression patterns and binding affinities to NKG2D. J Immunol. 2008;180(3):1678–85.

[67]. Sharma N, Flaherty K, Lezgiyeva K, Wagner DE, Klein AM, Ginty DD. The emergence of transcriptional identity in somatosensory neurons. Nature. 2020;577(7790):392–8.

[68]. Jung M, Dourado M, Maksymetz J, Jacobson A, Laufer BI, Baca M, et al. Cross-species transcriptomic atlas of dorsal root ganglia reveals species-specific programs for sensory function. Nat Commun. 2023;14(1):366.

[69]. Usoskin D, Furlan A, Islam S, Abdo H, Lonnerberg P, Lou D, et al. Unbiased classification of sensory neuron types by large-scale single-cell RNA sequencing. Nat Neurosci. 2015;18(1):145–53.

[70]. Krauter D, Kupari J, Usoskin D, Su J, Hu Y, Zhang M-D, et al. Spatial organization, chromatin accessibility and gene-regulatory programs defining mouse sensory neurons. Commun Biol. 2025:(in press).

[71]. Marmigere F, Ernfors P. Specification and connectivity of neuronal subtypes in the sensory lineage. Nature reviews Neuroscience. 2007;8(2):114–27.

[72]. Goswami SC, Mishra SK, Maric D, Kaszas K, Gonnella GL, Clokie SJ, et al. Molecular signatures of mouse TRPV1-lineage neurons revealed by RNA-Seq transcriptome analysis. The journal of pain : official journal of the American Pain Society. 2014;15(12):1338–59.

[73]. Hammond DL, Ackerman L, Holdsworth R, Elzey B. Effects of spinal nerve ligation on immunohistochemically identified neurons in the L4 and L5 dorsal root ganglia of the rat. J Comp Neurol. 2004;475(4):575–89.

[74]. Liu CN, Wall PD, Ben-Dor E, Michaelis M, Amir R, Devor M. Tactile allodynia in the absence of C-fiber activation: altered firing properties of DRG neurons following spinal nerve injury. Pain. 2000;85(3):503–21.

[75]. Cavanaugh DJ, Lee H, Lo L, Shields SD, Zylka MJ, Basbaum AI, et al. Distinct subsets of unmyelinated primary sensory fibers mediate behavioral responses to noxious thermal and mechanical stimuli. Proceedings of the National Academy of Sciences of the United States of America. 2009;106(22):9075–80.

[76]. Brumovsky P, Villar MJ, Hokfelt T. Tyrosine hydroxylase is expressed in a subpopulation of small dorsal root ganglion neurons in the adult mouse. Exp Neurol. 2006;200(1):153–65.

[77]. Mishra SK, Tisel SM, Orestes P, Bhangoo SK, Hoon MA. TRPV1-lineage neurons are required for thermal sensation. EMBO J. 2011;30(3):582–93.

[78]. Taylor-Clark TE, Wu KY, Thompson JA, Yang K, Bahia PK, Ajmo JM. Thy1.2 YFP-16 transgenic mouse labels a subset of large-diameter sensory neurons that lack TRPV1 expression. PLoS One. 2015;10(3):e0119538.

[79]. Rebuffet L, Melsen JE, Escaliere B, Basurto-Lozada D, Bhandoola A, Bjorkstrom NK, et al. High-dimensional single-cell analysis of human natural killer cell heterogeneity. Nat Immunol. 2024;25(8):1474–88.

[80]. Heming M, Wolbert J, Börsch A-L, Thomas C, Mausberg AK, Lu IN, et al. Multi-omic characterization of human sural nerves across polyneuropathies. bioRxiv. 2024:2024.12.05.627043.

[81]. Davies AJ, Lleixa C, Siles AM, Gourlay DS, Berridge G, Dejnirattisai W, et al. Guillain-Barre Syndrome Following Zika Virus Infection Is Associated With a Diverse Spectrum of Peripheral Nerve Reactive Antibodies. Neurol Neuroimmunol Neuroinflamm. 2023;10(1).

[82]. Kalia AK, Rosseler C, Granja-Vazquez R, Ahmad A, Pancrazio JJ, Neureiter A, et al. How to differentiate induced pluripotent stem cells into sensory neurons for disease modelling: a functional assessment. Stem Cell Res Ther. 2024;15(1):99.

[83]. Carayannopoulos LN, Naidenko OV, Kinder J, Ho EL, Fremont DH, Yokoyama W. Ligands for murine NKG2D display heterogeneous binding behavior. Eur J Immunol. 2002;32(3):597–605.

[84]. Djelloul M, Popa N, Pelletier F, Raguenez G, Boucraut J. RAE-1 expression is induced during experimental autoimmune encephalomyelitis and is correlated with microglia cell proliferation. Brain Behav Immun. 2016;58:209–17.

[85]. Hu G, Huang K, Hu Y, Du G, Xue Z, Zhu X, et al. Single-cell RNA-seq reveals distinct injury responses in different types of DRG sensory neurons. Sci Rep. 2016;6:31851.

[86]. Zeisel A, Hochgerner H, Lonnerberg P, Johnsson A, Memic F, van der Zwan J, et al. Molecular Architecture of the Mouse Nervous System. Cell. 2018;174(4):999–1014 e22.

[87]. Chen CL, Broom DC, Liu Y, de Nooij JC, Li Z, Cen C, et al. Runx1 determines nociceptive sensory neuron phenotype and is required for thermal and neuropathic pain. Neuron. 2006;49(3):365–77.

[88]. Shields SD, Ahn HS, Yang Y, Han C, Seal RP, Wood JN, et al. Nav1.8 expression is not restricted to nociceptors in mouse peripheral nervous system. Pain. 2012;153(10):2017–30.

[89]. Woodhall M, Mgbachi V, Fox H, Irani S, Waters P. Utility of Live Cell-Based Assays for Autoimmune Neurology Diagnostics. J Appl Lab Med. 2022;7(1):391–3.

[90]. Irani SR, Gelfand JM, Al-Diwani A, Vincent A. Cell-surface central nervous system autoantibodies: clinical relevance and emerging paradigms. Ann Neurol. 2014;76(2):168–84.

[91]. Davies AJ, Rinaldi S, Costigan M, Oh SB. Cytotoxic Immunity in Peripheral Nerve Injury and Pain. Front Neurosci. 2020;14:142.

[92]. Reynders A, Mantilleri A, Malapert P, Rialle S, Nidelet S, Laffray S, et al. Transcriptional Profiling of Cutaneous MRGPRD Free Nerve Endings and C-LTMRs. Cell Rep. 2015;10(6):1007–19.

[93]. Seal RP, Wang X, Guan Y, Raja SN, Woodbury CJ, Basbaum AI, et al. Injury-induced mechanical hypersensitivity requires C-low threshold mechanoreceptors. Nature. 2009;462(7273):651–5.

[94]. George DS, Jayaraj ND, Pacifico P, Ren D, Sriram N, Miller RE, et al. The Mas-related G protein-coupled receptor d (Mrgprd) mediates pain hypersensitivity in painful diabetic neuropathy. Pain. 2024;165(5):1154–68.

[95]. Warwick C, Cassidy C, Hachisuka J, Wright MC, Baumbauer KM, Adelman PC, et al. MrgprdCre lineage neurons mediate optogenetic allodynia through an emergent polysynaptic circuit. Pain. 2021;162(7):2120–31.

[96]. Cooper AH, Barry AM, Chrysostomidou P, Lolignier R, Wang J, Redondo Canales M, et al. Peripheral nerve injury results in a biased loss of sensory neuron subpopulations. Pain. 2024;165(12):2863–76.

[97]. Gangadharan V, Zheng H, Taberner FJ, Landry J, Nees TA, Pistolic J, et al. Neuropathic pain caused by miswiring and abnormal end organ targeting. Nature. 2022;606(7912):137–45.

[98]. Feng J, Zhao Y, Xie Z, Zang K, Sviben S, Hu X, et al. Miswiring of Merkel cell and pruriceptive C fiber drives the itch-scratch cycle. Sci Transl Med. 2022;14(653):eabn4819.

[99]. Jeon SM, Pradeep A, Chang D, McDonough L, Chen Y, Latremoliere A, et al. Skin Reinnervation by Collateral Sprouting Following Spared Nerve Injury in Mice. The Journal of neuroscience : the official journal of the Society for Neuroscience. 2024;44(15).

[100]. Kim HW, Shim SW, Zhao AM, Roh D, Han HM, Middleton SJ, et al. Long-term tactile hypersensitivity after nerve crush injury in mice is characterized by the persistence of intact sensory axons. Pain. 2023;164(10):2327–42.

[101]. Jung H, Hsiung B, Pestal K, Procyk E, Raulet DH. RAE-1 ligands for the NKG2D receptor are regulated by E2F transcription factors, which control cell cycle entry. J Exp Med. 2012;209(13):2409–22.

[102]. Greene TT, Tokuyama M, Knudsen GM, Kunz M, Lin J, Greninger AL, et al. A Herpesviral induction of RAE-1 NKG2D ligand expression occurs through release of HDAC mediated repression. Elife. 2016;5.

[103]. Gasser S, Orsulic S, Brown EJ, Raulet DH. The DNA damage pathway regulates innate immune system ligands of the NKG2D receptor. Nature. 2005;436(7054):1186–90.

[104]. Routes JM, Ryan S, Morris K, Takaki R, Cerwenka A, Lanier LL. Adenovirus serotype 5 E1A sensitizes tumor cells to NKG2D-dependent NK cell lysis and tumor rejection. J Exp Med. 2005;202(11):1477–82.

[105]. Nomura M, Zou Z, Joh T, Takihara Y, Matsuda Y, Shimada K. Genomic structures and characterization of Rae1 family members encoding GPI-anchored cell surface proteins and expressed predominantly in embryonic mouse brain. J Biochem. 1996;120(5):987–95.

[106]. Popa N, Cedile O, Pollet-Villard X, Bagnis C, Durbec P, Boucraut J. RAE-1 is expressed in the adult subventricular zone and controls cell proliferation of neurospheres. Glia. 2011;59(1):35–44.

[107]. Gan Y, Liu Q, Wu W, Yin JX, Bai XF, Shen R, et al. Ischemic neurons recruit natural killer cells that accelerate brain infarction. Proceedings of the National Academy of Sciences of the United States of America. 2014;111(7):2704–9.

[108]. Garofalo S, Cocozza G, Porzia A, Inghilleri M, Raspa M, Scavizzi F, et al. Natural killer cells modulate motor neuron-immune cell cross talk in models of Amyotrophic Lateral Sclerosis. Nat Commun. 2020;11(1):1773.

[109]. Donovan LJ, Brewer CL, Bond SF, Laslavic AM, Pena Lopez A, Colman L, et al. Aging and injury drive neuronal senescence in the dorsal root ganglia. Nat Neurosci. 2025;28(5):985–97.

[110]. Techameena P, Feng X, Zhang K, Hadjab S. The single-cell transcriptomic atlas iPain identifies senescence of nociceptors as a therapeutical target for chronic pain treatment. Nat Commun. 2024;15(1):8585.

[111]. Ogasawara K, Hamerman JA, Ehrlich LR, Bour-Jordan H, Santamaria P, Bluestone JA, et al. NKG2D blockade prevents autoimmune diabetes in NOD mice. Immunity. 2004;20(6):757–67.

[112]. Markiewicz MA, Wise EL, Buchwald ZS, Pinto AK, Zafirova B, Polic B, et al. RAE1epsilon ligand expressed on pancreatic islets recruits NKG2D receptor-expressing cytotoxic T cells independent of T cell receptor recognition. Immunity. 2012;36(1):132–41.

[113]. Gravett AM, Dalgleish AG, Copier J. In vitro culture with gemcitabine augments death receptor and NKG2D ligand expression on tumour cells. Sci Rep. 2019;9(1):1544.

[114]. Niu C, Jin H, Li M, Zhu S, Zhou L, Jin F, et al. Low-dose bortezomib increases the expression of NKG2D and DNAM-1 ligands and enhances induced NK and gammadelta T cell-mediated lysis in multiple myeloma. Oncotarget. 2017;8(4):5954–64.

[115]. Iannello A, Raulet DH. Immune surveillance of unhealthy cells by natural killer cells. Cold Spring Harb Symp Quant Biol. 2013;78:249–57.

[116]. Yang D, Sun B, Li S, Wei W, Liu X, Cui X, et al. NKG2D-CAR T cells eliminate senescent cells in aged mice and nonhuman primates. Sci Transl Med. 2023;15(709):eadd1951.

[117]. Udit S, Blake K, Chiu IM. Somatosensory and autonomic neuronal regulation of the immune response. Nature reviews Neuroscience. 2022;23(3):157–71.

[118]. Verma V, Drury GL, Parisien M, Ozdag Acarli AN, Al-Aubodah TA, Nijnik A, et al. Unbiased immune profiling reveals a natural killer cell-peripheral nerve axis in fibromyalgia. Pain. 2022;163(7):e821–e36.

[119]. Shiers S, Klein RM, Price TJ. Quantitative differences in neuronal subpopulations between mouse and human dorsal root ganglia demonstrated with RNAscope in situ hybridization. Pain. 2020;161(10):2410–24.

[120]. Shiers SI, Sankaranarayanan I, Jeevakumar V, Cervantes A, Reese JC, Price TJ. Convergence of peptidergic and non-peptidergic protein markers in the human dorsal root ganglion and spinal dorsal horn. J Comp Neurol. 2021;529(10):2771–88.

[121]. Middleton SJ, Barry AM, Comini M, Li Y, Ray PR, Shiers S, et al. Studying human nociceptors: from fundamentals to clinic. Brain. 2021;144(5):1312–35.

[122]. Graus F, Campo E, Cruz-Sanchez F, Ribalta T, Palacin A. Expression of lymphocyte, macrophage and class I and II major histocompatibility complex antigens in normal human dorsal root ganglia. J Neurol Sci. 1990;98(2-3):203–11.

[123]. Mathieson S, Maher CG, Terwee CB, Folly de Campos T, Lin CW. Neuropathic pain screening questionnaires have limited measurement properties. A systematic review. J Clin Epidemiol. 2015;68(8):957–66.

[124]. Finnerup NB, Haroutounian S, Kamerman P, Baron R, Bennett DLH, Bouhassira D, et al. Neuropathic pain: an updated grading system for research and clinical practice. Pain. 2016;157(8):1599–606.

[125]. Tracey I, Woolf CJ, Andrews NA. Composite Pain Biomarker Signatures for Objective Assessment and Effective Treatment. Neuron. 2019;101(5):783–800.

[126]. Fillingim M, Tanguay-Sabourin C, Parisien M, Zare A, Guglietti GV, Norman J, et al. Biological markers and psychosocial factors predict chronic pain conditions. Nat Hum Behav. 2025.

[127]. Fundaun J, Kolski M, Molina-Alvarez M, Baskozos G, Schmid AB. Types and Concentrations of Blood-Based Biomarkers in Adults With Peripheral Neuropathies: A Systematic Review and Meta-analysis. JAMA Netw Open. 2022;5(12):e2248593.

[128]. Denk F, Ramer LM, Erskine EL, Nassar MA, Bogdanov Y, Signore M, et al. Tamoxifen induces cellular stress in the nervous system by inhibiting cholesterol synthesis. Acta Neuropathol Commun. 2015;3:74.

[129]. Olson JA, Leveson-Gower DB, Gill S, Baker J, Beilhack A, Negrin RS. NK cells mediate reduction of GVHD by inhibiting activated, alloreactive T cells while retaining GVT effects. Blood. 2010;115(21):4293–301.

[130]. Ruggeri L, Capanni M, Casucci M, Volpi I, Tosti A, Perruccio K, et al. Role of natural killer cell alloreactivity in HLA-mismatched hematopoietic stem cell transplantation. Blood. 1999;94(1):333–9.

[131]. Wang S, Davies AJ. (2025) Flow cytometry data - Purity and stimulation of human NK cells. (Zenodo). 10.5281/zenodo.15603183 (Accessed July 2025)

